# The genetics of cortical organisation and development: a study of 2,347 neuroimaging phenotypes

**DOI:** 10.1101/2022.09.08.507084

**Authors:** Varun Warrier, Eva-Maria Stauffer, Qin Qin Huang, Emilie M. Wigdor, Eric A.W. Slob, Jakob Seidlitz, Lisa Ronan, Sofie Valk, Travis T. Mallard, Andrew D. Grotzinger, Rafael Romero-Garcia, Simon Baron-Cohen, Daniel H. Geschwind, Madeline Lancaster, Graham K. Murray, Michael J. Gandal, Aaron Alexander-Bloch, Hyejung Won, Hilary C. Martin, Edward T. Bullmore, Richard A.I. Bethlehem

## Abstract

Our understanding of the genetic architecture of the human cerebral cortex is limited both in terms of the diversity of brain structural phenotypes and the anatomical granularity of their associations with genetic variants. Here, we conducted genome-wide association meta-analysis of 13 structural and diffusion magnetic resonance imaging derived cortical phenotypes, measured globally and at 180 bilaterally averaged regions in 36,843 individuals from the UK Biobank and the ABCD cohorts. These phenotypes include cortical thickness, surface area, grey matter volume, and measures of folding, neurite density, and water diffusion. We identified 4,349 experiment-wide significant loci associated with global and regional phenotypes. Multiple lines of analyses identified four genetic latent structures and causal relationships between surface area and some measures of cortical folding. These latent structures partly relate to different underlying gene expression trajectories during development and are enriched for different cell types. We also identified differential enrichment for neurodevelopmental and constrained genes and demonstrate that common genetic variants associated with surface area and volume specifically are associated with cephalic disorders. Finally, we identified complex inter-phenotype and inter-regional genetic relationships among the 13 phenotypes which reflect developmental differences among them. These analyses help refine the role of common genetic variants in human cortical development and organisation.

**One sentence summary:** GWAS of 2,347 neuroimaging phenotypes shed light on the global and regional genetic organisation of the cortex, underlying cellular and developmental processes, and links to neurodevelopmental and cephalic disorders.

## Main

The human cerebral cortex is morphologically complex, with extensive inter-individual and inter-regional variation associated with cognition, behaviour, health, development and ageing (*1*–*4*). This variation is partly genetic (*5*–*8*), with several loci identified primarily (although not exclusively) with cortical thickness, surface area, and volume (*6*, *9*–*12*). Less is known about the common variant genetics (including single nucleotide polymorphisms or SNPs) associated with more complex cortical morphometric phenotypes, such as gyrification or curvature or with microstructural MRI measures of cortical myelination and cyto-architecture. Consequently, the extent of shared genetics across surface area, cortical thickness, volume and the aforementioned MRI phenotypes is unknown. These relatively under-investigated cortical phenotypes may be as important as thickness, volume or other more traditional cortical phenotypes in determining individual differences in cognition, behaviour, and health (*13*–*15*). Second, we still do not fully understand how complex cellular and molecular mechanisms of neurodevelopment give rise to these distinct cortical brain phenotypes. Mapping this may help us better pinpoint genetic mechanisms underlying structural cortical abnormalities. Third, linked to this, the degree to which genes that are constrained (i.e., genes which are intolerant to damaging variants) or genes linked to neurodevelopmental conditions are enriched for common genetic variants for cortical morphology is unknown. Fourth, it is also unclear if common genetic variants contribute to cephalic disorders, although the impact of *de novo* damaging variants has been well documented (*16*). Finally, the role of common genetic variants in regional cortical phenotypes and organisation is also unclear. This is important as regional organisation may partly emerge from heterochronous regional differences in gene expression (*17*).

To address these questions, we conducted 2,347 genome-wide association studies (GWAS) (for 13 global and 2,334 regional phenotypes that met quality control [see **Methods**]) of cortical brain morphology in 36,843 individuals from the UK Biobank (UKB) (*19*) and the Adolescent Brain Cognitive Development (ABCD) (*21*) cohorts. These included eight cortical macrostructural phenotypes extracted from high resolution anatomical MRI sequences, and five cortical microstructural phenotypes extracted from diffusion MRI sequences, which were estimated both globally and across 180 bilaterally averaged regions based on the Human Connectome Project parcellation scheme (*18*). The 13 phenotypes included: cortical thickness (CT), cortical surface area (SA), grey matter volume (volume), folding index (FI), intrinsic curvature index (ICI), local gyrification index (LGI), mean curvature (MC), gaussian curvature (GC), fractional anisotropy (FA), mean diffusivity (MD), intracellular volume fraction (ICVF, also called neurite density index or NDI), isotropic volume fraction (ISOVF), and orientation dispersion index (ODI) (**Figure 1, Methods**). We share this GWAS summary statistics openly.

**Figure 1:**
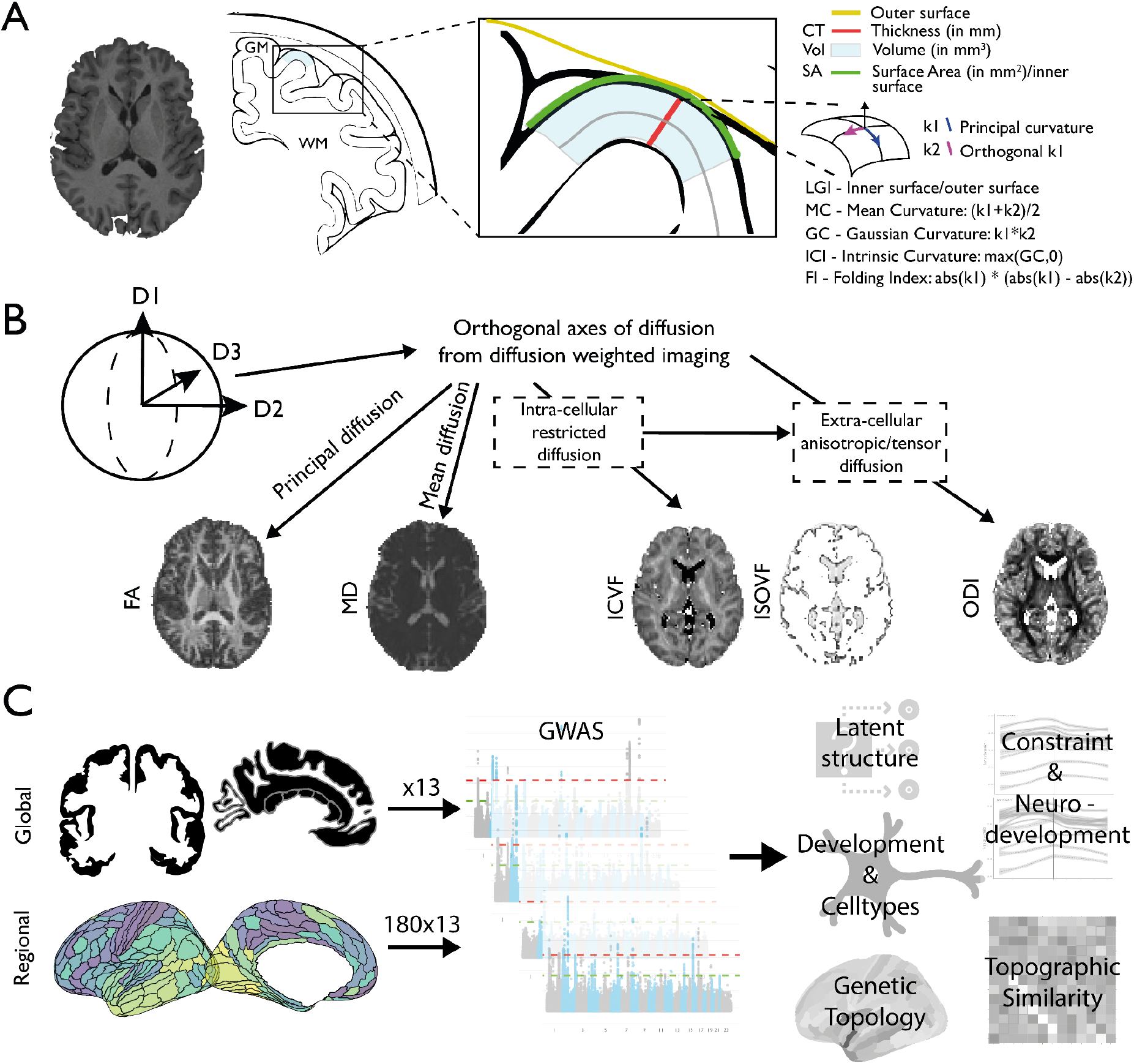
Schematic overview of 13 brain MRI phenotypes and the genetic analyses. (A) We considered eight cortical macrostructural phenotypes: cortical thickness (CT), cortical surface area (SA), grey matter volume (volume), folding index (FI), intrinsic curvature index (ICI), local gyrification index (LGI), mean curvature (MC), and Gaussian curvature (GC). (B) We also considered five cortical microstructural phenotypes: fractional anisotropy (FA), mean diffusivity (MD), intracellular volume fraction (ICVF, also called Neurite Density Index [NDI]), isotropic volume fraction (ISOVF), and orientation dispersion index (ODI). (C) Each phenotype was measured globally (total or mean for the whole cortex) and regionally at each of 180 bilaterally averaged cortical regions defined by the Human Connectome Project parcellation scheme. We conducted genome-wide association studies of all phenotypes after removing outliers, and investigated latent structure of all phenotypes, developmental trajectories, and cell type specificity and genetic organisation.

### Genome-wide associations of global cortical phenotypes

We first focused on the common variant genetics of 13 global structural MRI cortical phenotypes, calculated from the total or average of all 180 cortical areas (henceforth “global phenotypes”, **Figure 1, Methods**). We conducted GWAS for the 13 global phenotypes in the UKB after rigorous quality control and restricting the sample to participants of predominantly European genetic ancestries^1^ (N_max_ = 31,977). We identified 314 approximately independent (r^2^ < 0.1, 1000 kb) genome-wide significant (p < 5 × 10^−8^) loci for the global phenotypes. 81 of these were significant at the more stringent experiment-wide significance threshold (**Supplementary Table (ST) 1, Methods**). We additionally conducted GWAS for the same 13 global phenotypes in individuals of predominantly European genetic ancestries in ABCD (N_max_ = 4,866). For 237 GWAS loci in UKB for which data were available in ABCD, 204 SNPs (86%) had concordant sign of genetic association (p < 0.001, two-tailed binomial sign test) with modest positive correlation of effect size (*r*=0.54, 95% confidence interval [CI] 0.45-0.63; **Supplementary Figure (SF) 1**). 75 (31%) of these SNPs had p-values (p)<0.05 in ABCD with concordant effect direction. Additionally, Linkage disequilibrium score (LDSC) based (*22*) genetic correlations between UKB and ABCD were positive and high (genetic correlations ranged from 0.2 to 1, **SF 1, ST 2**) for all 13 phenotypes except MD, albeit with wide confidence intervals due to the relatively small size of the ABCD dataset. The high genetic correlation given the age difference between UKB (45-81 yrs at scanning) and ABCD (9-10 yrs at scanning) is notable as brain structure and its genetic influences change over time (*23*, *24*).

Given the observed shared genetics between UKB and ABCD, we conducted inverse-variance weighted meta-analyses (*25*) to combine the GWAS results across both UKB and ABCD. These meta-analyses identified 367 genome-wide significant loci, of which 89 were significant at an experiment-wide threshold **(ST 3)**. This ranged from 50 genome-wide significant loci (18 experiment-wide significant) for SA to 6 GWAS loci (with 0 experiment-wide significant) for FA (**Figure 2**), with some SNPs being associated with two or more phenotypes. In total there were 75 independent experiment-wide significant SNPs across all phenotypes. For all GWAS, the attenuation ratio (**Methods)**was not statistically different from 0 (**ST 4**), indicating no inflation in test statistics due to uncontrolled population stratification. All phenotypes had significant SNP heritabilities (LDSC (*26*): 0.06 for FA to 0.37 for SA), with higher SNP heritabilities for cortical macrostructural metrics (**ST 4**) compared to cortical microstructural phenotypes.

**Figure 2:**
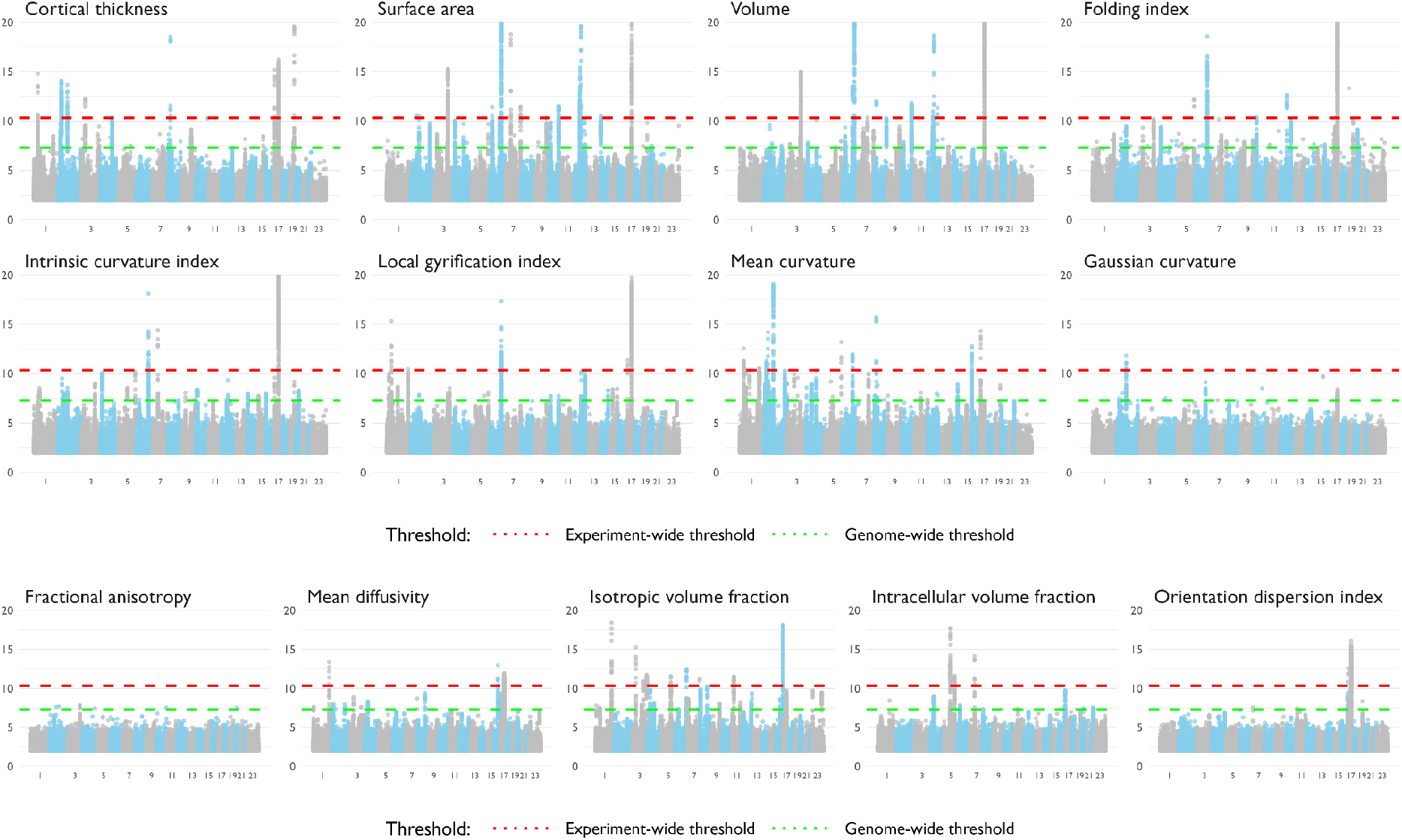
Manhattan plots of GWAS meta-analysis of 13 global MRI phenotypes. Green dotted line indicates the threshold for genome-wide significance (p = 5×10^−8^), and the red dotted line indicates the threshold for experiment-wide significance (p = 4.58×10^−11^). Each dot on the × axis indicates a SNP, and the y axis indicates the negative log10 p-value.

For SA and CT, we identified high genetic correlations with a larger GWAS based on a partly overlapping sample from the ENIGMA consortium (SA: r_g_ = 0.91 ±0.03; CT: r_g_ = 0.83 ±0.04) (*6*). Notably, despite the smaller sample size of the current GWAS meta-analyses, we identified a higher number of genome-wide significant loci for both SA (50 vs 19) and CT (31 vs 3), and had higher statistical power measured using mean χ^2^ (SA: 1.30 for current GWAS vs 1.23 for ENIGMA; CT: 1.23 for current GWAS vs 1.18 for ENIGMA). The gain in power is likely due to reduced heterogeneity in imaging and genotyping in the current study compared to ENIGMA. All three significant loci for CT and 15 out of the 19 significant loci for SA from ENIGMA were significant in our GWAS with concordant effect directions.

With the exception of CT, SA and volume, GWAS have not been conducted for the other cortical phenotypes. We investigated if any of the 75 independent experiment-wide significant SNPs identified in our GWAS were associated (i.e, p < 5×10^−8^) with any other neuroimaging phenotype using four databases (**Methods**). We found that 17 of these 75 SNPs were not associated with any other neuroimaging phenotype suggesting that these are novel associations **(ST 5)**.

### Four clusters explain shared genetics among most of the cortical phenotypes

Previous research has identified both distinct (e.g., between SA and CT) and shared genetics between some (e.g., SA and volume) cortical phenotypes (*6*, *27*). It seems likely that some genetic effects are pleiotropic for multiple cortical phenotypes, but it is unclear how these phenotypes cluster based on shared genetics. To address this question, we estimated bivariate genetic correlations and bivariate phenotypic correlations across all 13 global phenotypes (**ST 6; Figure 3A**). The overall pattern of correlations between measures was highly similar between the genotypic and phenotypic correlation matrices (Mantel’s test, r=0.89, P=1×10^−4^), in line with Cheverud’s conjecture that phenotypic correlations are representative of genetic correlations (*28*). We specifically identified high positive (genetic and phenotypic) correlations between five macrostructural phenotypes - SA, volume, LGI, ICI, and FI. Clustering of the genetic correlation matrix using multiple different methods consistently found that 12 of the 13 phenotypes (excluding only CT) formed four clusters relating to cortical expansion, curvature, water diffusion, and neurite density and orientation (**SF 2**). For the phenotypic correlation matrix, 11 of the 13 phenotypes formed four clusters, with CT and ICVF clustering separately (**SF 2**). Overall we identified high similarity between phenotypic and genetic cluster dendrograms (r_cophenetic_ = 0.76 (Ward D linkage) to 0.86 (Complete linkage)).

**Fig 3.**
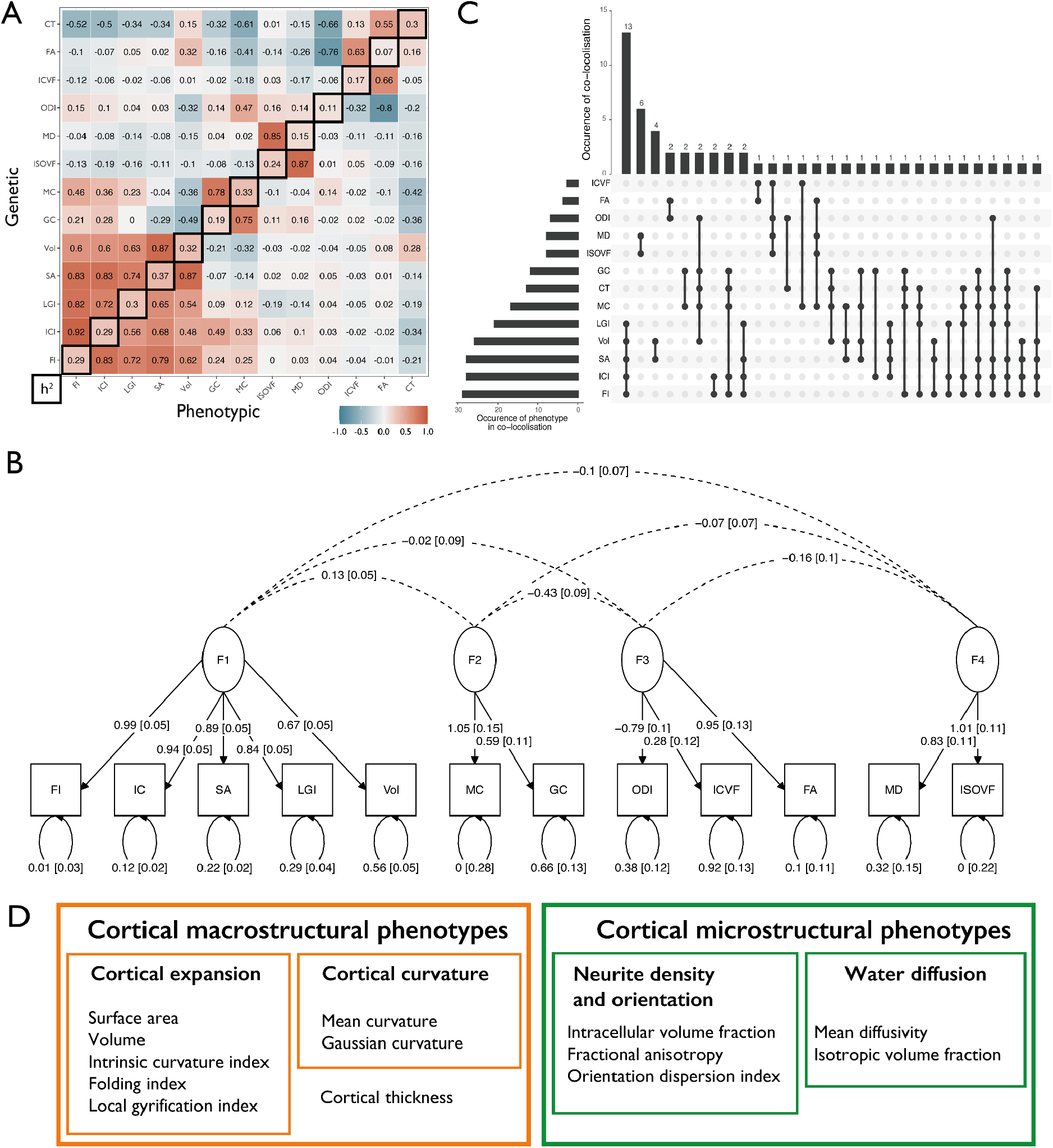
Pleiotropy among the 13 global phenotypes demonstrated by genetic/phenotypic correlations, structural equation modelling and co-localisation analysis. A: Phenotypic and genetic correlation matrices. The upper matrix triangle shows bivariate genetic correlations for each pair of phenotypes estimated using LDSC, the lower triangle shows the pairwise phenotypic correlations (Spearman’s coefficient). The diagonal indicates the SNP heritability of each phenotype based on LDSC. Phenotypes are ordered based on hierarchical clustering of the genetic bivariate correlation (hierarchical clustering on the phenotypic correlation matrix resulted in a near identical ordering). B: Genomic SEM path diagram demonstrating the underlying latent structure of 12 of the 13 global phenotypes and the interfactor genetic correlations. Covariance relationships = dotted arrows connecting two variables, variance estimates = double-headed arrows connecting variable to itself, regression relationships = one-headed arrows pointing from independent variable to dependent variable. Circles indicate latent variables, squares indicate measured phenotypes. The model was identified using unit variance identification such that the variance of the latent factors was set to one, and the dotted arrows across the factors can be interpreted as genetic correlation estimates. C: UpSet plot of the results of co-localisation analysis demonstrating the numbers of genomic loci that co-localise between the 13 phenotypes. The dots correspond to the co-localised clusters, with the number of clusters in the vertical bars. The number of times a phenotype co-localises is provided in the horizontal bars. D: Summary of clusters identified through GSEM and colocalisation analyses and their relationship with other terms used in this study. Abbreviations: cortical thickness (CT), cortical surface area (SA), grey matter volume (Vol), folding index (FI), intrinsic curvature index (ICI), local gyrification index (LGI), mean curvature (MC), and Gaussian curvature (GC), fractional anisotropy (FA), mean diffusivity (MD), intracellular volume fraction (ICVF), isotropic volume fraction (ISOVF), and orientation dispersion index (ODI).

To formalise this exploratory analysis of pleiotropic effects on distinct subsets of global brain phenotypes, we used genomic structural equation modelling (SEM) (*29*) to identify latent structures among the 13 global phenotypes. After excluding CT due to singleton-clustering and moderate genetic correlations (rg between −0.3 and −0.7 with eight of the 12 cortical phenotypes (**Figure 3A**; **SF 2**)), exploratory followed by confirmatory factor analyses identified a correlated four-factor model with acceptable fit (CFI = 0.89, SRMR = 0.13; **ST7, Figure 3B**). The four factors were similar to the four clusters and relate to cortical expansion (factor 1), curvature (factor 2), neurite density and orientation (factor 3), and water diffusion (factor 4). Phenotypic factor analyses produced four similar factors, albeit only after the removal of CT which did not cluster with any phenotypes and ICI which exhibited high cross-loading onto two factors (**Supplementary Note 1 [SN], SF 3**).

Co-localisation analysis of the experiment-wide significant associations supported the clustering and genomic-SEM analyses. Co-localisation analysis, which tests if genetic variants at a genetic locus are shared between two or more phenotypes, identified 56 co-localised genetic clusters among the global phenotypes (posterior probability of co-localisation> 0.6). The highest number of co-localised loci was for cortical expansion phenotypes, followed by water diffusion, neurite density and orientation phenotypes, and then curvature (**ST 8, Figure 3C**).

Cluster analysis, genomic-SEM, and co-localisation analysis thus convergently indicate four latent factors, each phenotypically represented by two or more MRI phenotypes. However, there is considerable shared genetics between the latent factors. Furthermore, CT was not included in the latent trait analyses. Given this, we conduct all downstream analysis at the level of individual phenotypes, but for consistency, additionally interpret results through the lens of four latent phenotypes and using the terms macrostructural and microstructural phenotypes (**Figure 3D**).

### Causal relationships between cortical expansion phenotypes

The genetic correlations between the phenotypes can be due to pleiotropy or causality. Consequently, we employed Mendelian randomisation (MR) (*30*) to investigate whether the genetic relationships between phenotypes represent causal mechanisms, especially among the five cortical expansion phenotypes. We tested three theories of causation, and corrected for multiple testing. First, consistent with the Radial Unit Hypothesis (*31*) which suggests that surface area emerges from the number of cortical columns but thickness emerges from the number of cells within a cortical column, we would not expect causal effects between SA and CT. Indeed, we observed no significant evidence for causal association between SA and CT. Second, since volume is geometrically related to SA and estimated by the product of SA and CT, we expected to find a bidirectional causal relationship between SA and volume, and indeed, we found evidence for this. Third, previous research (*32*–*35*) suggests that sulco-gyral folding emerges from differential tangential expansion of the cortex, partly due to the heterogeneous cortical distribution of progenitor cells (*36*, *37*), suggesting a causal relationship of SA on folding (FI, LGI and ICI). Consistent with this, we found robust evidence that genetically predicted SA is associated with an increase in certain measures of folding (FI, LGI, and ICI); but no evidence for reciprocally robust causal effects of folding metrics on SA (**ST 9 - 11, SN 2, SF 4-5**). Together, these analyses suggest causal relationships between SA and some measures of folding.

### Global cortical phenotypes show differential developmental and cellular associations

The complex genetic architecture among the 13 global phenotypes likely represents shared and distinct developmental and cellular processes. To better understand this, we first aggregated SNP-based p-values to gene-based p-values using two methods (MAGMA (*38*) and H-MAGMA (*39*)), and then investigated if these genes exhibited specific developmental trajectories of gene expression using post-mortem brain tissue data from PsychEncode (*40*). We excluded FA due to the small number of genes identified. Genes associated with six of the seven macrostructural phenotypes had high relative expression prenatally, a peak in the late mid-gestation period (~19-22 post-conception weeks), and a decline in gene expression postnatally (**Figure 4A**). In contrast, the four microstructural phenotypes were associated with genes that had peak expression at birth, followed by a less steep decline, or increased expression postnatally. Inspection of the phenotypes based on the four previous clusters did not identify any additional obvious trends in gene-expression trajectories. Linear mixed effects models confirmed significantly higher prenatal gene expression for most cortical macrostructural phenotypes, and higher postnatal gene expression for most microstructural phenotypes (**ST 12**).

**Fig 4.**
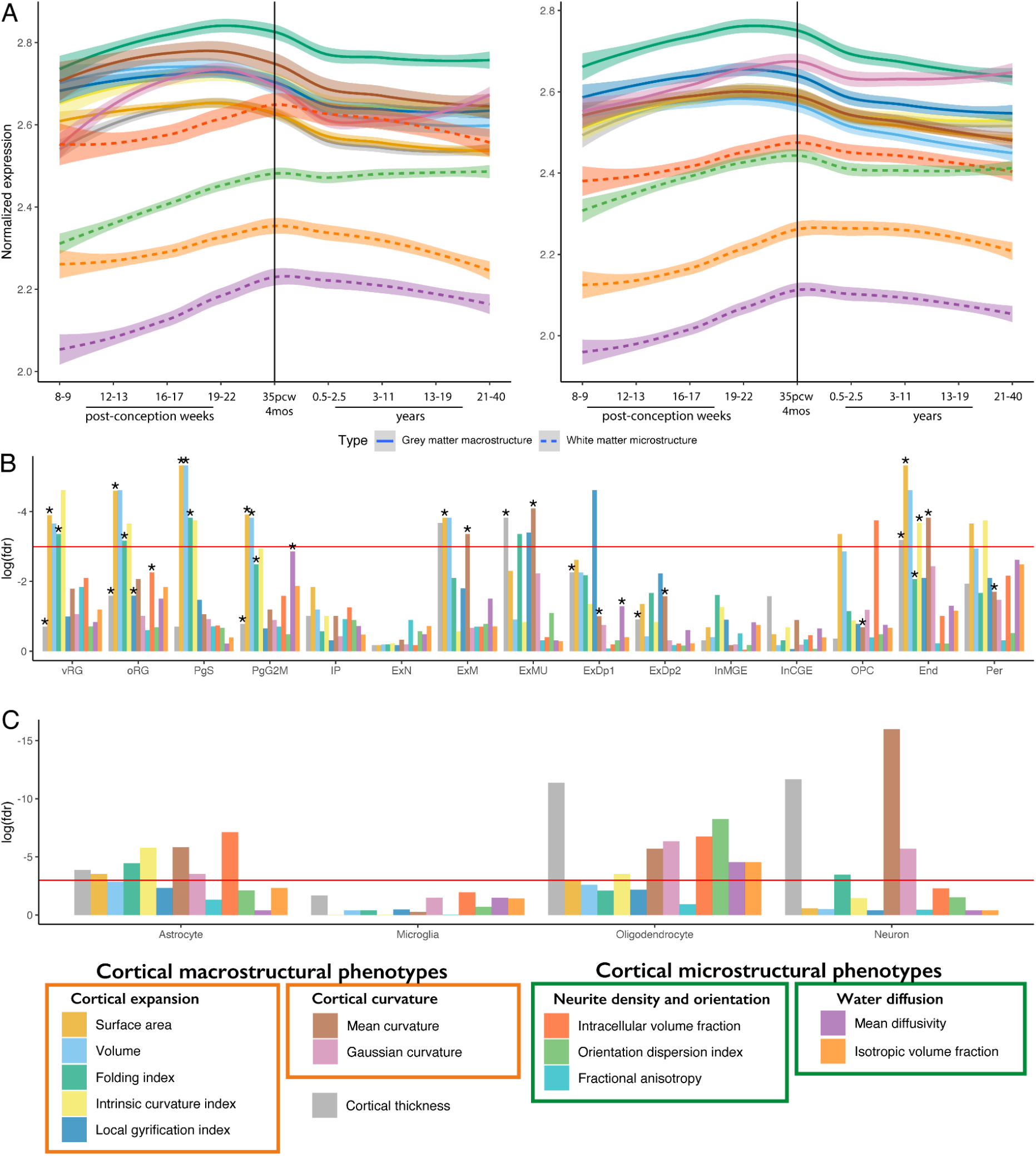
Enrichment of GWAS signals in different cell-types during development. A: Developmental trajectories of average gene expression in cortical postmortem-bulk RNA data (PsychEncode) for all significant genes (FDR < 0.05) identified using HMAGMA (left panel) or MAGMA (right panel) for 12 of the 13 global phenotypes. Data for FA not shown as too few genes were identified as significant. B: Results of enrichment analyses for cell-specific gene expression from mid gestation. FDR corrected log10 p-values for gene enrichment using genes identified from MAGMA are plotted. Additionally, significant enrichments identified using HMAGMA genes are indicated with an asterisk C: Results of enrichment analyses from cell-specific epigenetic marks from postnatal cortex identified using LDSC-based enrichment. Cell types for panel B: vRG = ventral radial glia, oRG = outer radial glia, PgS and PgG2M = cycling progenitors, S phase and G2-M phase respectively, IP = intermediate progenitors, ExN = Migrating excitatory neurons, ExM = Maturing Excitatory neurons, ExMU = Maturing excitatory neuron, upper enriched, ExDp1 = Excitatory deep layer neurons 1, ExDp2 = Excitatory deep layer neurons 2, InMGE = MGE Interneuron, InCGE = CGE interneuron, OPC = oligodendrocyte precursor cells, End = endothelial cells, Per = pericytes.

The differences in developmental trajectories likely reflect different underlying cellular compositions for these phenotypes. We used data from both single-cell RNA seq (scRNA-seq, enrichment using HMAGMA and MAGMA genes) and epigenetic data (LDSC-based (*26*) enrichment) to identify candidate cell types. Focusing on the developing brain, using sc-RNAseq data from psychENCODE (*17*), we identified enrichment for intermediate progenitor cells for SA, volume, and FI. (**ST 13**). To provide further temporal resolution into the developing brain, we investigated enrichment using scRNA-seq data from the first trimester (6-10 post conception weeks [pcw]) (*41*) and scRNA-seq (*42*) and scATAC-seq (*43*) data from mid-gestation (marked by neural progenitor expansion) (*44*–*46*). We did not identify any enrichment with cell types in the first trimester (**ST 14**) but FI, volume, and SA (cortical expansion phenotypes) were enriched for progenitor cells during mid-gestation (**ST 15 - 16, Figure 4B**), specifically for progenitor cells in the S-phase and G2-M phases of mitosis.

Additionally, CT and MC were enriched for multiple neuronal and glial cell types in both datasets, suggesting that these phenotypes are a composite of multiple cell types.

Considering the postnatal brain, there was no significant enrichment of genes in scRNA-seq data from psychENCODE (**ST 17**). However, analyses using epigenetic signatures of four broad cell types (*47*) (neurons, astrocytes, oligodendrocytes, microglia: **ST 18**) identified enrichment across multiple phenotypes (**Figure 4C**). For instance, cortical microstructural phenotypes were primarily enriched for epigenetic marks in oligodendrocytes and astrocytes, but not neurons, consistent with the idea that these phenotypes primarily reflect myelination and related processes. Taken together, these results demonstrate that genes underlying the 13 global phenotypes have different developmental trajectories reflecting specific cellular developmental dynamics.

### Cortical expansion phenotypes are associated with neurodevelopmental conditions

Given the enrichment of several of the global phenotypes with prenatal cellular and developmental processes, we hypothesised that these phenotypes are under negative selection pressure wherein damaging alleles are removed from the population by natural selection. So, we modelled the relationship between the minor allele frequency of the SNP and variance in effect size to quantify genome-wide signatures of selection using SBayesS (*48*). This suggested that the majority of the cortical macrostructural phenotypes are under significant negative selection (**Figure 5A**). The selection pressure for cortical microstructural phenotypes was weaker, and with the exception of ISOVF, not statistically significant (**ST 19**).

**Figure 5:**
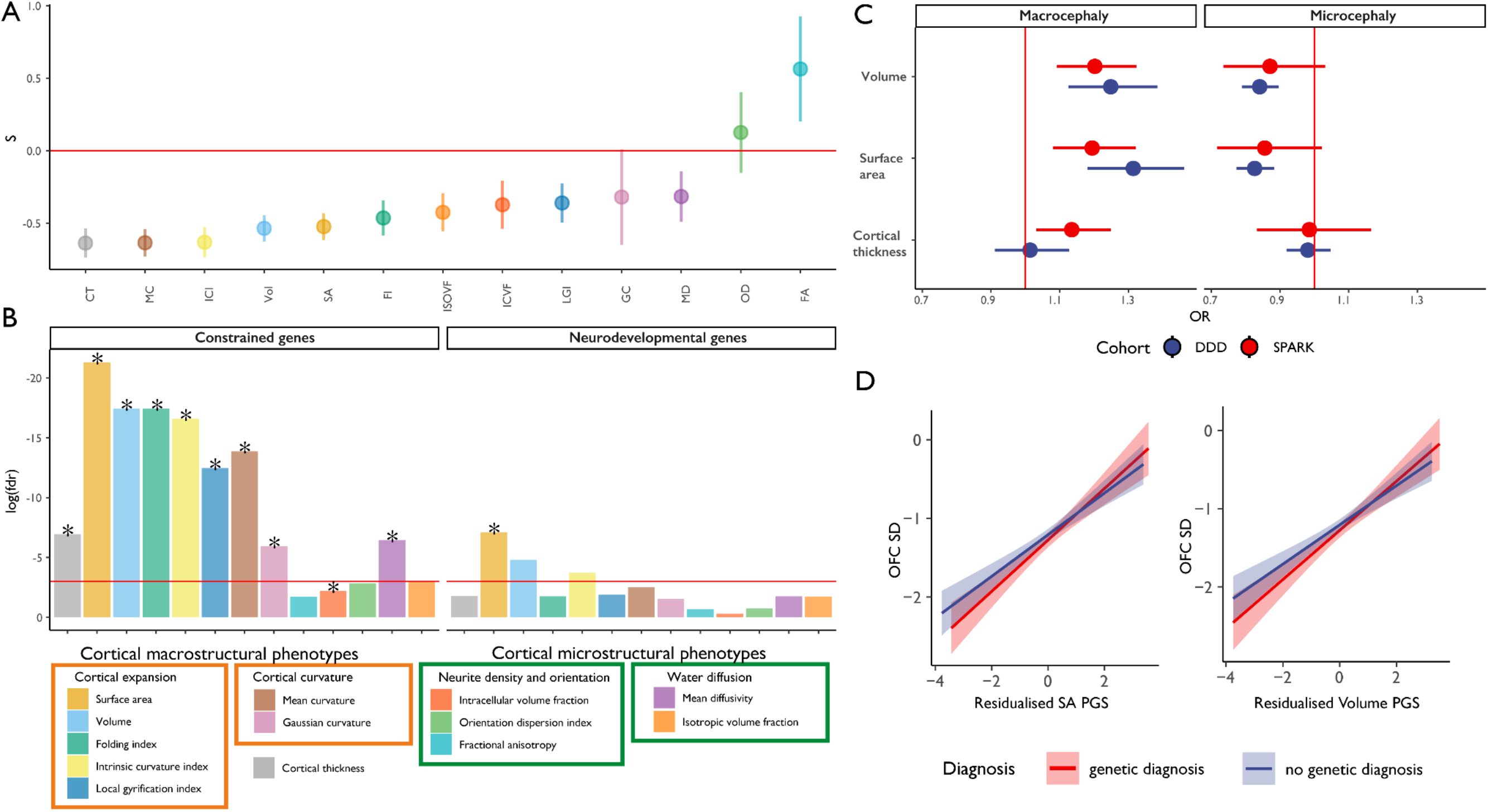
Signatures of constraint and links to neurodevelopment for the global phenotypes. A: Estimates of selection for the 13 cortical phenotypes. Selection coefficients (S) provided on the y axis (points). Bars indicate standard deviations for the selection coefficients. Negative values indicate lower-MAF alleles tend to have larger effect sizes B: Results of the enrichment analyses for constrained genes and genes associated with neurodevelopmental disorders using genes identified from MAGMA.FDR corrected log10 p-values for gene enrichment using genes identified from MAGMA are plotted. Additionally, significant enrichments identified using HMAGMA genes are indicated with an asterisk C: Odds ratio (OR) and 95% confidence intervals for macrocephaly and microcephaly compared to individuals with neither for 1 standard deviation increase in polygenic scores for volume, surface area, and cortical thickness in the DDD and SPARK cohorts. D: Line plots demonstrating the linear association between genetic principal component corrected polygenic scores (surface area (SA) and volume) and standardised (compared to general population) occipital-frontal circumference (OFC SD) for individuals with or without a genetic diagnosis in the DDD cohort.

We further reasoned that GWAS signals for some of these phenotypes would be enriched for constrained genes (i.e., genes from which damaging variants are removed by natural selection) (*49*), genes associated with severe neurodevelopmental conditions (*50*), or genes related to microcephaly. Indeed, cortical macrostructural phenotypes were significantly enriched for highly constrained genes (pLOUEF < 0.37) and, SA was enriched for genes associated with neurodevelopmental conditions (**ST 20, Figure 5B**) using both MAGMA and HMAGMA based enrichment. However, we identified no enrichment for genes linked to microcephaly, possibly because (1) several genes associated with microcephaly and other relevant cephalic disorders (e.g., lissencephaly and holoprosencephaly) are yet to be discovered or properly documented, or (2) clinical microcephaly (and macrocephaly) might be genetically distinct from normative variation in brain size.

To investigate the second hypothesis, we tested the effect of polygenic scores for SA, volume and CT on risk of cephalic conditions in a sample of 6,916 individuals with severe neurodevelopmental disorders from the DDD study (*51*, *52*). Polygenic scores for SA and volume, but not CT were associated with both macrocephaly and microcephaly in the expected directions (**Figure 5C, ST 21**). We obtained consistent results in another cohort (SPARK) (*53*) consisting of autistic individuals and their siblings, some of whom have macrocephaly or microcephaly (N=25,621). Furthermore, in the DDD cohort, polygenic scores for both volume and SA were significantly associated with occipital-frontal circumference standardised for age and sex, in both individuals with and without a genetic diagnosis (**Figure 5D**). These results demonstrate that SNPs associated with cortical expansion phenotypes are enriched in constrained genes and genes linked to neurodevelopmental conditions, and contribute to cephalic disorders and quantitative variation in occipital-frontal circumference even among individuals with severe neurodevelopmental disorders.

### Prioritising candidate genes supports progenitor proliferation in cortical expansion

Given the previous enrichment and polygenic association with neurodevelopmental and cephalic disorders, we were interested in identifying potential causal genes from the GWAS and investigate if these genes are associated with cephalic or neurodevelopmental conditions. Whilst HMAGMA and MAGMA are useful methods to aggregate SNP based information to provide gene-based p-values, they do not necessarily identify either the causal variant or the candidate gene. We thus conducted functionally informed fine-mapping of all experiment-wide significant loci using Polyfun (*54*) to identify causal variants. For 29 of these loci we were able to finemap to fewer than five credible variants, and for eight, a single credible variant (**ST 22)**. We used nine overlapping methods to identify candidate genes (**Methods**) and identified 181 candidate genes associated with one of the 13 global phenotypes (**ST 23**) by at least one method. From this list, we defined prioritised candidate genes if they were supported by at least two experimental methods, leading to 40 different prioritised candidate genes, including 19 in the 17q21.31 region (**ST 24**). Of these, 29 were identified for cortical expansion phenotypes, four for curvature phenotypes, 13 for neurite density and orientation phenotypes, 14 for water diffusion phenotypes, and 12 for CT, with considerable overlap between the phenotypic domains.

Several genes identified for cortical expansion phenotypes are involved in mitosis, neural progenitor proliferation, and cephalic and neurodevelopmental conditions including *ATR (55)*, *CENPW (56)*, *KANSL1 (57)* and *HMGA2 (58–60)*. Mutations in *ATR* cause Seckel syndrome, characterised by dwarfism, severe microcephaly, and intellectual disability *(55)*. *KANSL1* is associated with Koolen-de Vries syndrome, characterised by global developmental delays, and with over 50% of published individuals having microcephaly *(57)*. Mutations in *HMGA2* lead to macrocephaly and Silver-Russell syndrome (*60*). The overlap between fine-mapped genes from common variants and genes implicated through rare variants suggest convergence between rare and common variants. The genes identified for the cortical expansion phenotypes were enriched for the wnt signalling pathway (GO:1904953, q = 0.04) which regulates progenitor proliferation and cortical size (*61*).

Some genes implicated in CT and neurite density and orientation phenotypes involved in axogenesis and neuronal migration including *VCAN (62)* and *MACF1*, mutations in which cause lissencephaly and defects in neuronal migration and axon guidance (*63*). Finally, genes associated with water diffusion phenotypes included *MOBP*, which encodes a structural component of the myelin sheath, the neuronal proline and glycine transporter gene *SLC6A20*, and the lipid gated potassium channel gene *KCNK2*.

### Genetic loci associated with regional cortical phenotypes

To identify genetic influences on regional measures of the 13 neuroimaging modalities, we conducted 2,338 GWAS using regional phenotypes measured for 180 bilaterally averaged regions of the cortex using the Human Connectome Parcellation scheme (*18*). We excluded ICI and FI in three regions because of a lack of variance. We did not adjust for global phenotypes as we were interested in identifying genetic loci in the global phenotypes that co-localised with regional phenotypes and because controlling for heritable, highly correlated covariates (|r| > ~0.5) biases the GWAS and increases the number of false positives (*64*) (**SN 3 and SF 13-14**). This additionally limits the utility of the GWAS summary statistics for downstream analyses, for example due to collider bias in MR (*64*, *65*). In total we identified 4,260 experiment-wide significant loci. The highest number were associated with regional SA (1,033) (**ST 25**). These loci were more likely to contain constrained regions of the genome (*66*) (p = 3.97×10^−3^, one-sided Wilcoxon rank-sum test). This enrichment was driven by loci that were significant for regional cortical expansion phenotypes (p = 4.38×10^−4^, one-sided Wilcoxon rank-sum test). The 4,263 loci clustered into 456 semi-independent regions when accounting for LD (r^2^ > 0.1, 1000 kb) agnostic of the neuroimaging phenotype, indicating widespread pleiotropy across the regional measures.

To understand the extent to which these signals reflect genetic influences on the global phenotypes, we used the “GWAS-by-subtraction” method to regress out a latent factor representing genetic variance (*67*) on global phenotypes for 3,216 of the experiment-wide significant loci (**Methods, ST 26**). 1,633 (50%) of these loci remained experiment-wide significant, and 3,049 (95%) remained genome-wide significant suggesting that the vast majority of these loci had statistically significant regional effects. In contrast the global genetic latent trait reached experiment-wide significance for 966 of these loci (30%), and 1,499 (46%) reached genome-wide significance, suggesting that as many as half of these loci are also associated with the global genetic latent trait. But this could be partly by construct, as the global phenotypes in this study are simply the sum of the regional phenotypes. Genome-wide modelling of genetic influences on global and regional factors (*68*) is beyond the scope of the current study, but can be pursued using summary data made available here.

To further identify shared genetic loci across regional and global phenotypes, we conducted co-localisation analyses across all experiment-wide significant loci (regional and global) for each of the 13 phenotypes separately (**ST 27**). We identified between 409 (for SA) and 17 (for FA) co-localised clusters, where we use the term “cluster” to refer to a group of phenotypes within one of the 13 neuroimaging modalities that share causal variants in an LD-defined genomic region. The largest cluster was at chr12: 65559695 - 67181144 (12q14.3) comprising the global SA and 156 other regional SA GWAS. This region includes the aforementioned *HMGA2*, associated with Silver-Russell Syndrome. For all phenotypes except FA and MD, larger clusters were more likely to include hits in the global GWAS (p < 0.05, one-sided Wilcoxon rank sum test). However, there were some large clusters that comprised only regional GWAS, suggesting more localised regional effects. Visual inspection of all clusters with cluster size of 30+ GWAS revealed that topologically closer regions were more likely to have higher genetic co-localisation (**SF 6 - 7**). Furthermore, median geodesic distance between regions within a cluster was smaller than the median geodesic distance between regions within and outside a cluster (p < 2×10^−16^, wilcoxon rank sum test).

Clusters that included the global GWAS also exhibited broader regional patterns of co-localisation. For example, a locus at chr6:125424383-127540461 which includes *CENPW* is associated with FI and ICI both globally and in over 30 regions in the superior (dorsal) cortex (**SF 8**). *CENPW* exhibits regional differences in gene expression in the developing cortex (*69*). These analyses demonstrate that SNPs associated with global phenotypes may be associated with only some regional phenotypes.

As with the global features, regional cortical macrostructural phenotypes showed an on average higher heritability compared to regional cortical microstructural phenotypes (**SF 9, ST 28**) (t = −19.4, p < 2×10^−16^, F_(12,2327)_ = 420.7). There was however widespread regional variability in SNP heritability of cortical morphology (**SF 9**). Given this variability, we evaluated if SNP heritability systematically varied across previously established functional (Yeo and Krienen communities) (*70*) and morphological (Mesulam classes) parcellations (*71*) of the cortex. Permutation analyses that account for spatial correlation between regions (spin-permutation) (*72*) revealed that only CT had relatively higher heritability in idiotypic sensory areas (Mesulam), and a similar profile was observed for the sensory-motor network (Yeo and Krienen) (*70*) (**ST 29, SF 10**). This may reflect better histological and functional demarcation of the sensory-motor regions relative to other regions. Overall, these results suggest limited evidence of SNP heritability for cortical morphology being preferentially larger or smaller in known functional and morphological organisational classes.

### Differential regional and cross-morphological genetic organisation of the cortex

The high-resolution parcellation scheme used in this study also allows us to evaluate the protomap hypothesis, which suggests that regional differentiation of the cortex is intrinsically (genetically) determined early in cortical development (*31*, *73*). If this is true, then we would expect regions that are spatially closer to each other to be genetically more similar. Partly supporting this, genetic correlations were moderately correlated with geodesic distances among the 180 regions for each of the 13 phenotypes (r = 0.57 for LGI to 0.13 for ICVF, p = 0.001 for all tests, Mantel test, **ST 30**).

We further investigated if regional genetic correlations were higher within either functionally similar networks (Yeo and Krienen communities (*70*)) and morphologically similar classes of laminar differentiation (Mesulam classes (*71*)). Across multiple phenotypes we identified higher genetic correlations among Mesulam’s heteromodal association cortical regions but not in any of the Yeo-Krienen communities (*70*) (**ST 31, SF 11**).

To better understand if the 13 phenotypes are similar in their pattern of regional genetic correlations, we calculated cophenetic correlation coefficients amongst all 13 neuroimaging modalities using the regional genetic correlation matrices. Grouping based on cophenetic correlations identified four clusters with similar regional genetic correlation patterns (Cluster 1: SA, volume, and LGI; Cluster 2: all folding measures and CT; Cluster 3: FA and OD; and Cluster 4: MD and ISOVF, **Figure 6A**). Similar clusters were also observed when using regional phenotypic correlation matrices. These clusters differed from the clusters identified from the global phenotypes in that FI and ICI clustered together with CT and other measures of curvature. This suggests that clusters based on shared genetics of global phenotype moderately overlap with clusters based on regional genetic organisation.

**Fig 6.**
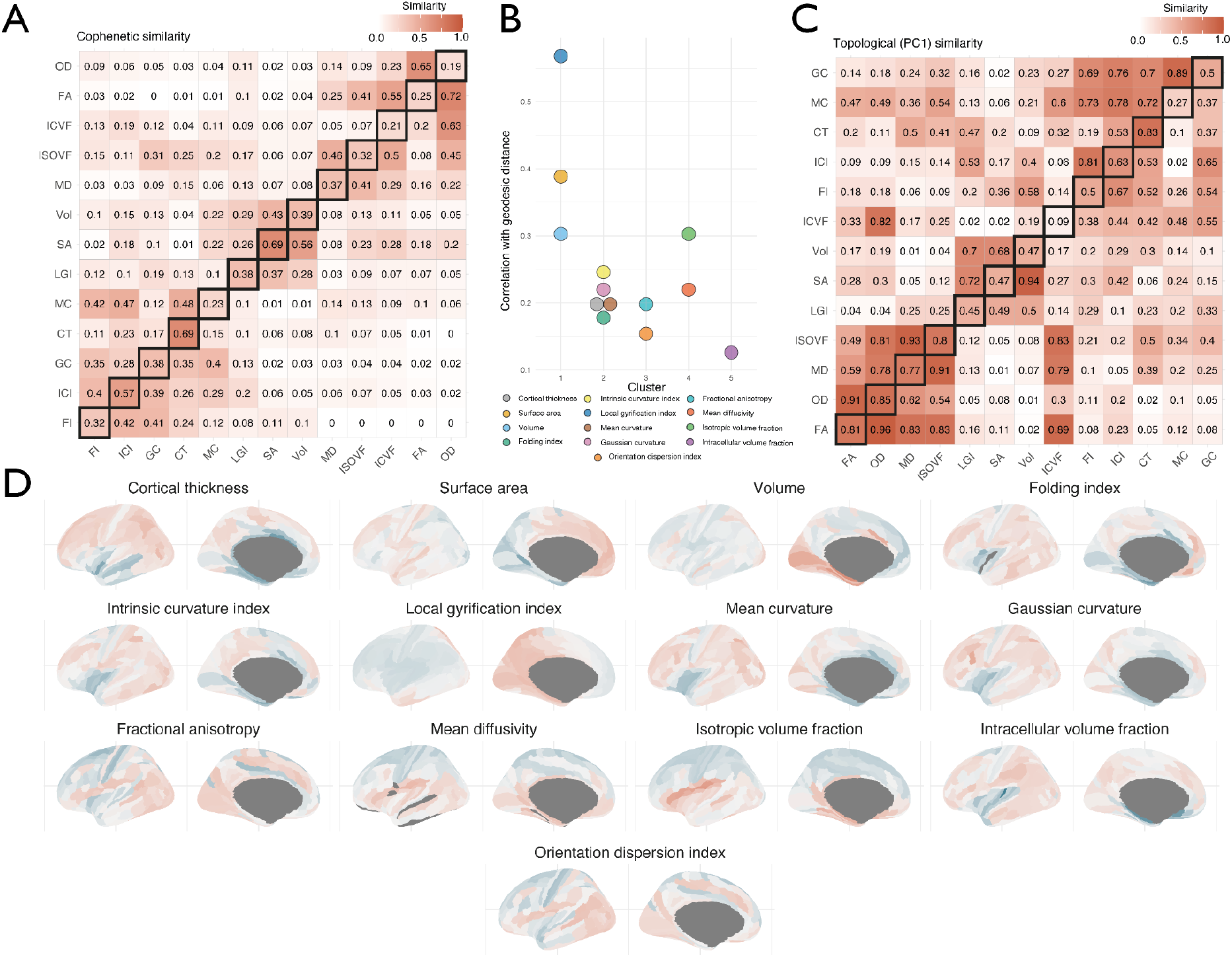
Topographic similarity and principal component structure of cortical phenotypes. A: Cophenetic similarity matrix depicting the similarity between the region*region similarity matrices. The upper triangle shows the cophenetic genetic similarity, the lower triangle shows the cophenetic phenotypic similarity and diagonals show the phenotype-genotype cophenetic similarity across features. B: Correlation between network topology and geodesic distance organised by hierarchical clustering of the cophenetic similarity. C: Spatial correlation between the first principal component of each regional similarity matrix. The upper triangle shows the genetic similarity, the lower triangle shows the phenotypic similarity and diagonals show the phenotype-genotype correlation across. D: Topology of the first genetic principal components, with colour depicting the relative PCA eigenvalues. The colour scale thus indicates to what extent regions show more homogenous similarity (i.e., regions with more similar colour have more similar covariance), but the actual sign and magnitude are relative within each phenotype.

These four clusters were also distinguishable based on their correlation between regional geodesic distances and genetic correlation (**Figure 6B**). Cluster 1 phenotypes, which relate to progenitor proliferation, had the highest correlation between genetic correlation and geodesic distance between regions. This was followed by Cluster 4 (MD and ISOVF: the water diffusion phenotypes), which both increase with age in adults (*74*, *75*). We speculate that this patterning might reflect the heterochronous cellular and developmental trajectories of these phenotypes: regional differences in gene expression in the cortex exhibit a cup-shaped pattern with high regional differences in midgestation that re-emerge during adolescence and increase in adulthood (*17*, *76*).

To explore further how the 13 phenotypes differed in their regional organisation, we extracted the first principal component from their respective genetic correlation matrices. The first principal component explained between 25% (LGI) to 62% (MD) of the variance. The clustering of the neuroimaging modalities based on the similarity of the first principal component of the region-to-region similarity was similar to the clustering based on the cophentic correlations of the same region-to-region similarity (**Figure 6C**), suggesting that the first principal component largely captures regional genetic organisation. Visual inspection of the first principal component identified four different axes of variation: anterior-posterior (SA, volume, and LGI: Cluster 1 phenotypes), inferior-superior (ISOVF, MD: Cluster 4 phenotypes), and a mix of primary-association and inferior superior (CT, GC, MC, FI, ICI: Cluster 2 phenotypes, and ICVF) (**Figure 6D**). For OD and FA (Cluster 3 phenotypes), we were unable to identify a clear topological axis of variation. These are in line with patterns of gene expression in the human cortex: anterior-posterior gradients during development, and primary-association gradients postnatally up until adolescence and early adulthood (*77*), and in the inferior-superior direction for water diffusion phenotypes, which are late-emerging (*78*, *79*). Using the first principal component derived from regional phenotypic correlation, we identified clear axes of variation for SA, Volume (anterior-posterior) and LGI, CT (inferior superior), but not for the other phenotypes (**SF 12**). This likely reflects the additional influence of directionless non-genetic factors in the development of cortical microstructure and curvature.

## Discussion

Our results provide granular insights into the organisation and development of the human cortex and links to cephalic and neurodevelopmental conditions. We find that cortical macrostructural and microstructural phenotypes are genetically distinct, and are enriched for different cellular and developmental processes. These results suggest that SA, volume, and related measures of gyrification emerge from the earliest cellular processes in cortical development, related to progenitor proliferation. Cortical thickness and some measures of curvature are influenced by multiple, subsequent and sequential cellular processes, as indicated by widespread enrichment for different cell types. These early developmental processes are under purifying selection, and consequently, enriched for highly constrained genes and genes associated with neurodevelopmental conditions. Furthermore, even among individuals with severe developmental disorders (*51*, *52*), common genetic variants contribute to cephalic disorders, expanding our understanding of the role of common and rare genetic variants in developmental disorders. For the first time, we investigate causal relationships between cortical macrostructural phenotypes using Mendelian randomisation, and provide support for the differential tangential expansion hypothesis (*32*, *35*, *36*).

These differences between the phenotypes are also reflected in their regional organisation. Phenotypes that derive from early developmental processes and phenotypes that continue to develop during late adulthood show greatest shared genetics between topologically nearer regions, in line with findings from cortical gene expression (*77*). However, whilst this is in the anterior-posterior direction for the early emerging phenotypes, it is in the inferior-superior direction for the late emerging phenotypes. Notably, the anterior-posterior gradient is also observed using cell lineage-tracing in the brain (*80*). This suggests that cortical organisation is informed by distinct waves of molecular processes, some of which are highly directional.

Our analyses focused on individuals predominantly of European genetic ancestries, as we were limited by sample size, computational power, and methodology. With ongoing neuroimaging in the UK Biobank and other cohorts, these analyses should be expanded to include genetically diverse populations. Additionally, the current study investigates only the role of common genetic variants, and the extent to which rare genetic variants contribute to normative differences in cortical morphology is unclear. Finally, expanding the number of phenotypes such as fMRI, white-matter, and subcortical measures will provide a more precise atlas of the genetics of structure and function of the human brain, and the genetic relationships between them.

In conclusion, by conducting and analysing GWAS of 13 different neuroimaging modalities both globally and across 180 cortical regions we provide unprecedented insights into the genetic organisation, and development of the human cortex. We make this resource freely available to researchers for further analyses.

## Acknowledgements

VW is supported by St. Catharine’s College Cambridge. E-MS is supported by a Ph.D. studentship awarded by the Friends of Peterhouse. EAWS issupported by the NIHR Cambridge Biomedical Research Centre (BRC-1215-20014). The views expressed are those of the authors and not necessarily those of the NIHR or the Department of Health and Social Care. RAIB is supported by the Autism Research Trust. SBC received funding from the Wellcome Trust 214322\Z\18\Z. For the purpose of Open Access, the author has applied a CC BY public copyright licence to any Author Accepted Manuscript version arising from this submission. SBC also received funding from the Autism Centre of Excellence, SFARI, the Templeton World Charitable Fund, the MRC, and the NIHR Cambridge Biomedical Research Centre. The research was supported by the National Institute for Health Research (NIHR) Applied Research Collaboration East of England. JS was supported by NIMH T32MH019112-29 and K08MH120564. E.T.B. was supported by an NIHR Senior Investigator award and the Wellcome Trust collaborative award for the Neuroscience in Psychiatry Network. A.F.A.-B. was supported by NIMH K08MH120564. R.R.G was supported by EMERGIA Junta de Andalucía program (EMERGIA20_00139). S.L.V. was supported by Max Planck Gesellschaft, (Otto Hahn Award) and the Helmholtz Association’s Initiative and Networking Fund under the Helmholtz International Lab grant agreement InterLabs-0015, and the Canada First Research Excellence Fund (CFREF Competition 2, 2015–2016) awarded to the Healthy Brains, Healthy Lives initiative at McGill University, through the Helmholtz International BigBrain Analytics and Learning Laboratory (HIBALL). GKM was supported by MRC MR/W020025/1.

Data were curated and analysed using a computational facility funded by an MRC research infrastructure award (MR/M009041/1) to the School of Clinical Medicine, University of Cambridge and supported by the mental health theme of the NIHR Cambridge Biomedical Research Centre. The views expressed are those of the authors and not necessarily those of the NIH, NHS, the NIHR or the Department of Health and Social Care. Data used in the preparation of this article were obtained from the Adolescent Brain Cognitive DevelopmentSM (ABCD) Study (https://abcdstudy.org), held in the NIMH Data Archive (NDA). This is a multisite, longitudinal study designed to recruit more than 10,000 children age 9-10 and follow them over 10 years into early adulthood.

The ABCD Study^®^ is supported by the National Institutes of Health and additional federal partners under award numbers U01DA041048, U01DA050989, U01DA051016, U01DA041022, U01DA051018, U01DA051037, U01DA050987, U01DA041174, U01DA041106, U01DA041117, U01DA041028, U01DA041134, U01DA050988, U01DA051039, U01DA041156, U01DA041025, U01DA041120, U01DA051038, U01DA041148, U01DA041093, U01DA041089, U24DA041123, U24DA041147. A full list of supporters is available at https://abcdstudy.org/federal-partners.html. A listing of participating sites and a complete listing of the study investigators can be found at https://abcdstudy.org/consortium_members/. ABCD consortium investigators designed and implemented the study and/or provided data but did not necessarily participate in the analysis or writing of this report. This manuscript reflects the views of the authors and may not reflect the opinions or views of the NIH or ABCD consortium investigators.The ABCD data repository grows and changes over time. The ABCD data used in this report came from [NIMH Data Archive Digital Object Identifier (10.15154/1503209)]. DOIs can be found at http://dx.doi.org/10.15154/1503209.

For the purpose of open access, the authors have applied a Creative Commons Attribution (CC BY) licence to any Author Accepted Manuscript version arising from this submission. We thank Luke Kweku Abraham and Jennifer Asimit for helpful discussions. The authors do not declare any competing interests.

## Supplementary materials and methods

### Datasets

#### UK BioBank

The UK Biobank is a prospective cohort of 500,000 individuals from the UK. Of these individuals, 100,000 will undergo brain scanning (*5*, *19*, *81*), with approximately 40,000 scans having been completed when the current study commenced. Participants were excluded from the MRI study on the basis of standard MRI safety criteria such as metal implants, recent surgery, or conditions problematic for scanning such as hearing problems, breathing problems, or claustrophobia. Ethical procedures for the UK Biobank are controlled by the Ethics and Guidance council (http://www.ukbiobank.ac.uk/ethics), and the study was conducted in accordance with the UK Biobank an Ethics and Governance Framework document (http://www.ukbiobank.ac.uk/wp-content/uploads/2011/05/EGF20082.pdf), with institutional review board approval by the North West Multi-center Research Ethics Committee.

#### ABCD

The Adolescent Brain Cognitive Development (ABCD) study is an ongoing study of childhood and adolescence (*82*). Participants from the general population were recruited from all over the United States across 21 sites by providing select schools with information packets to all families with 8-10 year old students. Ethical approval was obtained from multiple institutional review boards.

#### Image processing

Structural minimally processed T1 and T2-FLAIR weighted data was obtained from UK BioBank (application 20904) and the ABCD study (via the NIH Data Archive Repository). These images were preprocessed with FreeSurfer (v6.0.1) (*83*) using the T2-FLAIR weighted image to improve pial surface reconstruction when available. Recon-all reconstruction included bias field correction, registration to stereotaxic space, intensity normalisation, skull-stripping, and white matter segmentation. A triangular surface tessellation fitted a deformable mesh model onto the white matter volume, providing grey-white and pial surfaces with >160,000 corresponding vertices registered to fsaverage standard space. When no T2-FLAIR image was available FreeSurfer reconstruction was done using the T1 weighted image only. Given systematic variation related to the inclusion of T2-FLAIR (see supplements), this was included as a confound variable in downstream analyses. Following reconstruction, the Human Connectome Project (HCP) parcellation (*18*) was aligned to each individual FreeSurfer average image and parcellated were values extracted. Reconstruction quality was assessed using the Euler index (*84*) and included as a covariate in subsequent analyses.

Structural diffusion weighted imaging was obtained in processed form from UK BioBank and ABCD in similar fashion. Neurite Orientation Dispersion and Density Indices (NODDI) parameters were estimated using the Accelerated Microstructure Imaging via Convex Optimization (AMICO) (*85*) processing approach from the minimally processed diffusion images. The T1 aligned parcellation template was co-registered to the diffusion weighted image using fsl FLIRT and regional values for FA, MD and the three NODDI parameters were extracted using AFNI’s 3dROIstats function for all of the 360 cortical regions included in the Human Connectome Parcellation and averaged across hemisphere to reduce the number of regions to 180 bilateral regions. In total 13 different imaging derived phenotypes were extracted using this pipeline:

1. Total Surface Area of the cortex (SA)
2. Total volume of the cortex (volume)
3. Average thickness of the cortex (CT)

Measures of curvature: We calculated five measures of curvature. Assuming two principal curvatures (k1 and k2), we can define the five measures of curvature as follows.

4. Total mean curvature (MC) = (k1 + k2)/2. MC is typically thought to measure extrinsic curvature. In other words, this is not curvature that is intrinsic to the surface, but rather extrinsic to the surface.
5. Total gaussian curvature (GC) = k1*k2
6. Total intrinsic curvature index (ICI) = max(K,0). In other words, if GC is positive, ICI is positive. If GC is negative, ICI is 0.
7. Total Folding Index (FI) = ABS (K1) * (ABS (K1) – ABS (K2)).
8. Total Local Gyrification Index (LGI) (*86*): Gyrification index quantifies the amount of the curvature that is buried within the sulcal folds, and is a measure of gyrification. This is computed by calculating the ratio of the area between an outer smoother surface and an inner surface tightly wrapping the pial surface. As it is a ratio, it is a unitless measure.

The above properties measure primarily tissue macrostructure. To better understand cortical microstructure, we calculated five measures from the diffusion weighted images(*87*).Since conventional diffusion parameters such as fractional anisotropy (FA) and mean diffusivity (MD) alone are not specific to the underlying microstructure of axons and dendrites (referred to, collectively, as neurites) we also extracted measures of Neurite Orientation Density and Dispersion Imaging (NODDI) (*88*, *89*).

9. Fractional anisotropy (FA) (*87*): FA is thought to be a measure of microstructural integrity. Higher FA values are thought to indicate fiber tracts (i.e., greater anisotropy). FA would be higher in areas of greater neurite density due to less isotropic water diffusion.
10. Mean diffusivity (MD) (*87*): MD measures the degree of displacement (or diffusivity) of water. It can be a measure of membrane density and degree of myelination. Lower membrane density and greater myelination is thought to decrease MD.

We calculated the following three metrics using NODDI. NODDI assumes three microstructural environments for the diffusion of water - intracellular, extracellular, and CSF (*90*). The intracellular environment is anisotropic and water diffusion in this environment can be quantified using

11. Intracellular volume fraction (ICVF): Also referred to as neurite density index or NDI, this is a measure of density of neurites (axons and dendrites). Higher ICVF values indicate that a greater fraction of the tissue consists of neurites.
12. Orientation dispersion index (ODI): This measures the orientation and spatial variation of the neurite fibres. Zero indicates perfectly aligned straight fibres, and one for completely isotropic fibres. Thus, larger values of ODI represent highly dispersed neurites and smaller values represent highly aligned neurites.
13. Isotropic volume fraction (ISOVF): This is a measure of water diffusion, typically representing cerebrospinal fluid and ventricles in the cortex.

We note that all phenotypes were standardised. Mean CT was calculated as the average across the 180 bilaterally averaged cortical regions. Due to this standardisation, the standardised scores from the average and total values will be identical.

#### Genome-wide association analyses

##### Genetic quality control in the UK Biobank

We included only individuals of self-reported white European ethnicity, and from this group of individuals, excluded individuals who were above +/− 5 SD from the means of the first two genetic principal components, had a genotyping rate < 95%, whose genetic sex did not match their reported sex, or had excessive genetic heterozygosity. We used all genotyped and imputed SNPs in the UK Biobank that had a minor allele frequency > 0.1%, did not deviate from Hardy-Weinberg equilibrium (P > 1×10^−6^), had a genotyping rate of 95%, and, for imputed SNPs, had an imputation R^2^ > 0.4. After quality control, we retained a maximum of 31,977 participants and 15,916,802 SNPs. We did not conduct GWAS for individuals in other ethnic groups as there were fewer than 400 individuals with imaging and genetic data after quality control in each of the other ethnic groups, which is insufficient sample size for linear mixed effect models for GWAS.

##### Genetic quality control in ABCD

Prior to imputation, we filtered SNPs with missingness > 90%, and deviations from Hardy Weinberg Equilibrium (p < 1×10^−6^). We removed individuals with missingness > 5% and whose genetic sex did not match their reported sex. As HWE and heterozygosity are incorrectly calculated in populations with diverse genetic ancestries, these steps were conducted in relatively homogenous genetic ancestral groups identified using principal-component based clustering after combining the data with the 1000 Genomes phase 3 data (*91*). Principal components were calculated using GENESIS (*92*) after accounting for relatedness between samples as calculated using KING (*93*). To identify genetically homogeneous groups, we used the first five principal components to identify clusters in the 1000 Genomes data using UMAP, identifying 7 broad populations - Non-finnish Europeans, Finnish Europeans, Africans, Americans, East Asians, South Asian, and Bengali. Then, using the first five PCs from the ABCD dataset, we projected individuals onto the seven clusters, identifying broadly homogeneous populations. HWE based filtering (p < 1×10^−6^), and removing individuals with excess heterozygosity (+/− 3 SD) was then conducted. On this final cleaned data, relatedness was calculated using KING and PCs were calculated after accounting for relatedness, all at the level of individual population categories. The data was then merged, and phased (Eagle v 2.4) and imputed (Minimac4) using the TOPMED Imputation Server. From the imputed data, we removed SNPs with poor imputation (r^2^ < 0.4) and minor allele frequency < 0.1% (N = 14,495,763 SNPs). We restricted our analyses to individuals of primarily European ancestries (N = 4,866).

##### Genome-wide association analyses

In both the UKB and ABCD we followed the same procedures outlined below. We conducted whole brain and regional GWAS analyses for the 13 phenotypes mentioned in the “Image Processing” section. For each region, we averaged the values bilaterally, resulting in a total of 180 regional phenotypes per phenotype. For ICI and FI we excluded regions “52”, “PI”, “PHA2” because of no variance. In total, we conducted 2,347 GWAS using FastGWA (*94*). FastGWA can simultaneously account for both relatedness and subtle population stratification in the analyses.

All phenotypes were scaled to a mean of 0 and a standard deviation of 1. We removed individuals who scored above or below 5 SDs from the mean for all phenotypes, as these are most likely technical outliers. Furthermore, these outliers skew the phenotypic scores and cannot be used in fastGWA which can produce false positives at stringent p-value or for SNPs with low minor allele frequencies (*94*). Additionally, we visually inspected histograms of all phenotypes and further removed outliers above or below 5 median absolute deviations for phenotypes with substantial skew, primarily for MD and FI. Additionally, to ensure that the GWAS were not confounded by fine-scale population stratification, among the individuals of European ancestry identified in UKB or ABCD, we removed individuals who were above or below 5 SDs from the mean of the first two genetic principal components. For all GWAS, we included age, age^2^, sex, agexsex, age^2^xsex, imaging centre, first 40 genetic principal components, mean framewise displacement, maximum framewise dispacement, and Euler Index (*84*) as covariates. In addition, for structural MRI metrics derived from T1, we included the availability of T2 scans as covariates as this influenced the calculation of these metrics.

For the regional GWAS, we chose not to include the respective global phenotypes for three reasons. First, adjusting for heritable and highly correlated phenotypes biases the GWAS estimates (*64*, *65*). All global phenotypes were substantially heritable and highly correlated with the regional phenotypes (detailed in **SN 3**). Second, including highly correlated and heritable covariates may result in collider bias for downstream analyses such as Mendelian Randomisation (*95*). Given that we wish to make the summary statistics available for researchers to conduct other analyses, including global phenotypes as a covariate can restrict the scope of downstream analyses. Finally, we were specifically interested in identifying SNPs with effects across the cortex, which may not have been possible if we had adjusted for global phenotypes. We note, methods such as genomic-SEM (*29*), mtCOJO (*96*), and multivariable MR (*97*) all allow adjustment for global GWAS in downstream analyses. Here, we used genomic-SEM to regress out the genetic effects of the global phenotype for the majority of experiment-wide significant SNPs (definition of which is detailed below), to identify the fraction of SNPs that remained significant. We note that modelling of global vs. local genetic effects at a genome-wide level as conducted elsewhere (*68*) is beyond the scope of this study.

We meta-analysed results from the UK Biobank and ABCD using inverse-variance weighted meta-analyses in Plink v1.9 (*98*), excluding SNPs that were absent from the UKB, given the difference in sample sizes (and consequently, statistical power) between the UKB and ABCD.

##### Multiple testing correction

Using matrix decomposition (*20*), we estimated that there were 1,092 independent phenotypes. Thus, we used an experiment-wide threshold of 4.58×10^−11^ (5×10^−8^/1,092) to correct for the multiple tests conducted. To identify significant loci, we clumped the GWAS using an r^2^ threshold of 0.1 over 1000kb. We used LD information available from a random sample of 5,000 unrelated individuals from the UK Biobank who were included in the GWAS.

#### Genetic correlation and causal analyses

##### Genetic correlation, SNP heritability estimation, clustering, and genomic structural equation modelling

For the global phenotypes we used LDSC (*22*, *26*) to compute genetic correlations and SNP heritability for the meta-analysed GWAS statistics, using LD weights from the North West European populations. Intercepts were not constrained. Heritability and genetic correlation (among 180 regions per phenotype for all 13 phenotypes) of the regional GWAS were calculated using LDSC as incorporated within genomic-SEM (*29*). Additionally, for the global phenotypes in the UK-biobank alone, we conducted GCTA-GREML (*99*) based SNP heritability using genetic relationship matrix calculated using all imputed SNPs included in the GWAS, for 30,765 unrelated individuals with neuroimaging GWAS. We applied the same quality control and used the same covariates as for the GWAS.

For the global phenotypes, clustering of the phenotypic and genetic correlation matrices were conducted on the Euclidean distance. As the final hierarchical clustering is dependent on the clustering method used, we used three different clustering methods (Average, Ward D, and Complete Linkage) and visualised the different clusters obtained. Cophenetic correlations (in R, stats (version 3.6.2)) were obtained by comparing the phenotypic and genetic dendrograms produced by the different clustering methods.

Genomic structural equation modelling was conducted using genomic-SEM (*29*) using summary GWAS statistics of the global cortical phenotypes. We conducted exploratory factor analyses using the even chromosomes, identified factor models, and conducted confirmatory factor analyses using the odd chromosomes. The final model was selected after multiple iterations based on both fit indices and theoretical predictions. Fit indices and path diagrams are provided for models based on all chromosomes.

For the regional phenotypes, we conducted 1000 spin permutations (*72*) tests to investigate if SNP heritability of regions or genetic correlation among regions were higher in regions falling within functionally (*70*) or morphologically similar classes (*71*). Spin permutation accounts for spatial correspondence between regions and generates null models using random rotations across the spherical cortical surface (*72*).

We investigated if the genetic correlation among regions were correlated with topological geodesic distances among regions using Mantel test (within each phenotype separately). We investigated if clustering of regions based on genetic correlations were similar between phenotypes based on cophenetic correlation.

##### Phenotypic correlation and principal component analysis

Comparable to region specific genetic correlations, we also generated region to region phenotypic correlation matrices (“structural covariance”) for both UK Biobank and ABCD cohorts by taking the Pearson correlation across subjects on the scaled and filtered data. UK Biobank and ABCD were then combined into a single meta covariance matrix using the “psychmeta” package in R (*100*).

We extracted the first principal component from the regional genetic correlation matrix and regional phenotypic correlation matrix for each of the thirteen phenotypes separately. This principal component analysis was done using a singular value decomposition of the centred and scaled similarity matrix using the “stats” package in R.

##### Co-localisation

To identify co-localised genomic regions among the 13 global phenotypes, we used Hyprcoloc (*101*). Hyprcoloc is robust to participant overlap, and can conduct multi-trait co-localisation using hundreds of GWAS. We restricted our analyses to experiment-wide significant loci, and mapped these onto predefined approximately independent LD blocks in individuals of European ancestry (approximately 1.6 Mb on average) (*102*). We did not adjust for either participant or known correlation between phenotypes, as the method gives reasonable results comparable to adjusting for correlation between phenotypes. We used the branch and bound divisive clustering algorithm incorporated in Hyprcoloc to identify clusters of phenotypes that co-localise at any given locus. We used the default variant specific prior probabilities in Hyprcoloc (*101*): prior 1 (probability that a SNP is associated with a single trait) as 1×10^−4^, and prior c (prior probability that the SNP is associated with a second trait) as 0.02. We identified co-localised genomic regions if the genomic regional association probability was 0.6 or higher. We used the same pipeline to investigate co-localisation for 180 regional GWAS and the global GWAS for each of the 13 phenotypes conducted separately.

##### Mendelian Randomisation

To investigate the causal effects of SA on other cortical macrostructural phenotypes, we conducted Mendelian Randomisation analyses (*30*) using global phenotypes. To avoid bias due to participant overlap, we randomly divided the UK Biobank into two groups of individuals (Group A: N = 15,884 of which males = 7,455; Group B, N = 15,899, of which males = 7,500) and conducted GWAS analyses in each of the groups separately for the eight cortical macrostructural phenotypes using the same pipeline as detailed above. We generated instruments which consisted of SNPs with p < 5×10^−8^ in the exposure, with minor allele frequency > 1%, and which were near-independent (clumping r^2^ = 0.001 using a 1000 kb window using data from 5,000 unrelated individuals from the UK Biobank). Where fewer than five SNPs met this criteria, we relaxed the p-value threshold to p < 1×10^−6^. Using SA GWAS generated in Group A as the exposure and the GWAS for the remaining 6 phenotypes in Group B as the outcome, we conducted inverse-variance weighted bi-directional Mendelian Randomisation analyses. To account for pleiotropy, we additionally conducted the following sensitivity analyses: (1) median weighted MR (majority-valid (*103*)); (2) MR-Egger (accounts for pleiotropy) (*104*) ; (3) MR PRESSO (detects and excludes outliers in the instrument (*105*)). Additionally, (4) for the significant MR results, to further account for correlated (vertical) pleiotropy, we conducted MR analyses using CAUSE (*106*) using two instruments: one with of SNPs with p <5×10^−8^, and another at a more relaxed threshold of p < 0.001. We investigated heterogeneity in the instrument using Cochran’s Q, and investigated if the Egger intercept was significant. We investigated if the orientation of the causal direction was correct using Steiger analyses (*107*), and conducted additional sensitivity analyses after removing SNPs that did not have the correct causal orientation. Finally, we inspected the scatter plot, forest plot, and plots generated from leave-one-out analyses to identify if the results were driven by a subset of the SNPs. Analyses (1) and (2), and the sensitivity analyses were conducted using the R-package TwosampleMR (*108*).

We repeated all MR analyses except for CAUSE using instruments generated in the UKB as the exposure and ABCD as the outcome. However, this was quasi-bidirectional, in that in both directions, the exposure was instruments generated in the UKB and the outcome was SNPs in the ABCD. We did not conduct CAUSE in this instance due to sample size imbalance which reduces statistical power.

Given substantial pleiotropy between the phenotypes, we identified significant MR associations if: 1. The p-value was < 0.0035 (bonferroni corrected threshold) in both the within UKB and the UKB-ABCD analyses for IVW, MR-Presso, and weighted median; 2 MR-Egger was in the consistent direction to the IVW (MR-Egger has lower statistical power so we did not require it to be statistically significant); 3. If Steiger analyses identified incorrect causal orientation, criteria 1 and 2 were met after Steiger filtering; and 4. Results were significant when MR was conducted using CAUSE which accounts for correlated pleiotropy. Analyses were conducted using the two-sample MR package (version 0.5.6)(*108*). Power-calculations (*109*) were conducted assuming a standard deviation in the exposure results in a 0.33 unit standard deviation change in the outcome, which is a medium effect size.

#### Gene-based association and enrichment analyses

##### Gene based association

We used MAGMA (version 1. 10) (*38*) to conduct gene-based association testing based on physical location. MAGMA assigns SNPs to the nearest gene. In line with previous analyses, we expanded the window to 35kb upstream and 10kb downstream of the gene to capture regulatory regions (*110*). In addition, we used HMAGMA (*39*) (using MAGMA v 1.08) to identify genes based on Hi-C mapping. In contrast to MAGMA, HMAGMA is able to map SNPs to genes based on long-range interactions and can account for tissue specific regulatory effects. To map developmental trajectories, we used Hi-C data from postnatal and prenatal human cortex (*39*, *111*). Subsequently, for enrichment analyses, we used Hi-C data from the prenatal cortex given that the majority of the phenotypes were either enriched for gene expression in the prenatal cortex or did not differ in gene expression between prenatal and postnatal cortex, and because many processes investigated occurred prenatally.

##### Developmental trajectories

To identify patterns of gene expression across cortical prenatal and postnatal windows, we used data from PsychEncode (*17*). The data was divided into 9 developmental windows: Window 1. 8 - 9 post-conception weeks (PCW); Window 2. 12 - 13 PCW; Window 3. 16 – 17 PCW; Window 4. 19 - 22 PCW; Window 5. 35 PCW - 4 months; Window 6. 6 months to 2.5 years; Window 7. 3 - 11 years; Window 8. 13 - 19 years; Window 9. 21 - 40 years. Gene expression values were log base 2 transformed after adding a pseudocount and normalised. For 12 of the 13 phenotypes the transformed expression values of all genes with q < 0.05 were averaged for each developmental window and smoothed LOESS curves were plotted. The excluded phenotype was FA as HMAGMA and MAGMA identified 1 and 0 genes with q < 0.05 respectively.

##### Enrichment analyses

To investigate enrichment for cell types, signatures of genomic constraint, and gene sets associated with neurodevelopmental and cephalic disorders, we conducted the following analyses. Within each gene set, significant results were identified after correcting for all 13 phenotypes using Benjamini-Hochberg FDR correction (q < 0.05).

To identify cell types in the prenatal and postnatal cortex we conducted enrichment analyses using (1) single cell gene expression data from PsychENCODE (*17*) using prenatal (5pcw to 125 days) and postnatal gene expression. To provide additional temporal resolution we also conducted analyses using (2) single cell gene expression data that spanned early cortical development (6 - 10 pcw) (*41*) ; (3) single cell gene expression data spanning mid gestation period of cortical development (17 - 18 pcw) (*42*); (4) Single-cell epigenomic data (scATAC-seq) from the midgestation period of cortical development (*43*); and (5) Cell-type specific (fluorescent-activated nuclei sorting isolated) epigenomic (ATAC-seq and ChiP-seq) data from postnatal cortex (*47*). Analyses for datasets 1 - 3 were conducted using MAGMA gene-set enrichment using genes identified by MAGMA and HMAGMA. Following previously described methods (*110*), we filtered out genes with non-unique names and genes not expressed in any cell types. Gene expression values were log base 2 transformed after adding a pseudocount and normalised. Mean cell-type specific gene values were calculated, and this was divided by the mean expression of the gene in all cells to get relative cell-type expression. We then selected the top 10% of genes with the highest relative expression in each cell type to conduct enrichment analyses using MAGMA gene-set enrichment analyses (*38*). Significant cell types were identified if q < 0.05 in analyses using both HMAGMA and MAGMA identified genes. Analyses for datasets 4 and 5 were conducted using conditional partitioned heritability analyses in LDSC (i.e., enrichment for a cell type after conditioning on all other cell types and baseline annotations) (*112*, *113*).

We used the same gene-enrichment pipeline as above to investigate gene enrichment for genes that are constrained (pLOUEF < 0.37) (*49*), genes associated with neurodevelopmental disorders (*50*) (662 genes with FDR < 0.05), and genes associated with severe microcephaly obtained from the Genomics England Panel (244 genes, signed off on March 2, 2022: <u>Severe microcephaly (Version 2.304)</u> (genomicsengland.co.uk)). Signatures of selection were identified using SBayesS (*48*).

#### Polygenic score association analyses

##### Genetic quality control and polygenic score generation

Polygenic scores (PGS) for SA, CT, and volume were calculated using the meta-analysed GWAS in a dataset of individuals with severe developmental disorders (DDD study, N = 6,916) and autistic individuals and their families (SPARK dataset, N = 25,621) using PRScs (*114*). PRScs is a Bayesian algorithm that infers posterior effect sizes of SNPs using continuous shrinkage, and does not require defining p-value thresholds. Details of genetic quality control in the DDD cohort in individuals of predominantly European ancestries are provided elsewhere (*52*). The data were re-imputed using the TOPMed reference panel and variants with low imputation quality (minimac4 r^2^ < 0.8) were excluded. We kept common (minor allele frequency > 1%) SNPs that are also in HapMap3 to calculate the PGS using PRScs. Genetic-ancestry QC of the SPARK dataset was conducted similar to the ABCD dataset and as detailed elsewhere (*115*, *116*). We calculated PGS on individuals of predominantly European ancestries as identified by genetic principal components. All polygenic scores were standardised with a mean zero and a standard deviation of 1 for all analyses.

##### Defining phenotypes in DDD and SPARK

In the DDD study, we used HPO terms assigned by clinicians to define macrocephaly (N=396 with HPO term “HP:0040194”, “HP:0000256”, “HP:0004482”, “HP:0004481”, “HP:0004488”, or “HP:0005490”) and microcephaly (N=1,198 with HPO term “HP:0040195”, “HP:0000252”, “HP:0005484”, “HP:0004485”, “HP:0000253”, “HP:0011451”, or “HP:0040196”). We also analysed occipital frontal circumference data (N=6,146), which were calculated as standard deviations from the mean given the proband’s gestational age at birth, age at time of measurement and sex. In the SPARK dataset, information about macro- and microcephaly were obtained from parental/caregiver reports of medical diagnoses.

##### Statistical analyses

Linear or logistic mixed-effect regressions (random intercepts for family, in SPARK) were conducted using either: (1) PGS for volume; (2) PGS for SA and CT in a multiple regression framework. Primary analyses were conducted using logistic regression, separately for macrocephaly and microcephaly (coded as 1) compared to controls (i.e. individuals in the cohort without microcephaly or macrocephaly) (coded as 0). In the DDD, we also conducted linear regression using standardised occipital frontal circumference. Additionally, we conducted linear regression with macrocephaly coded as 1, microcephaly as −1 and no diagnosis as 0. In the DDD study, we included sex, genetic diagnosis, and the first 10 genetic principal components as covariates. Specifically, we considered probands to be “diagnosed” if they had at least one variant reported to DECIPHER that had been confirmed as pathogenic or likely pathogenic (C/LP) by a clinician, or that had been predicted as P/LP by a computational algorithm based on the American College of Medical Genetics criteria, as described in Wright et al. (*117*). In SPARK, age, sex, autism diagnosis, and the first 10 genetic principal components were included as covariates. Significant results were identified after Benjamini-Hochberg FDR correction (q < 0.05) across all models.

#### Fine-mapping, Summary Mendelian Randomisation, and prioritising candidate genes

For all exome-wide significant loci in the global GWAS (N = 90), we conducted functionally informed fine-mapping using Polyfun (*54*), using SuSiE (*118*) as the fine-mapping method and with up to 5 causal variants per locus, with each locus defined 500 kb upstream and downstream of the sentinel variant. In-sample LD was obtained from 5,000 unrelated individuals included in the GWAS from the UK Biobank. We used precomputed prior causal probabilities from the UK Biobank as provided in Polyfun.

To link the variants in the 95% credible sets to genes, we used Hi-C data from (1) the prenatal brain germinal zone (*111*), (2) the prenatal brain cortical plate (*111*), (3) neurons from postnatal cortex (*119*), and (4) glia from postnatal cortex (*119*). Additionally, we (5) used Ensembl Variant Effect Predictor (*120*) to identify genes containing damaging missense (deleterious in SIFT and/or damaging/probably damaging/possibly damaging in PolyPhen) and protein truncating variants from the list of the 95% credible sets.

To identify candidate genes using relevant eQTL and methylation data, we further conducted summary Mendelian randomisation (SMR) (*121*). SMR was conducted for all 13 phenotypes, using cis-eQTL data from postmortem (6) prenatal (*122*) and (7) postnatal brains (*123*), and additionally (8) methylation data from postnatal brains (*124*). Within each phenotype, we identified significant genes by using Bonferroni correction for the total number of genes tested. We excluded significant genes with evidence to indicate that the MR association results are due to pleiotropy using the HEIDI test (HEIDI p < 0.01) (*121*).

Finally, (9) we identified the closest gene to each sentinel variant (i.e., the SNP with the lowest p-value in each locus). Where the variant was intergenic, we included both the closest upstream and downstream gene. From these nine methods, we identify a list of prioritised candidate genes if they are supported by at least two methods. We conducted GO enrichment analyses to identify biological pathways enriched for the prioritised candidates genes.

## Data availability

All summary statistics for the GWAS meta-analyses will be made available upon publication without restriction on https://portal.ide-cam.org.uk/overview/483.

## Code availability

https://github.com/ucam-department-of-psychiatry/UKB

https://github.com/ucam-department-of-psychiatry/ABCD

## Software used

1. FastGWA: <u>GCTA | Yang Lab</u> (westlake.edu.cn)
2. Plink 1.9 and <u>Plink 1.2: PLINK 1.9</u> (cog-genomics.org)
3. genomic-SEM: <u>GitHub - GenomicSEM/GenomicSEM: R-package for structural equation modeling based on GWAS summary data</u>
4. LDSC v1.01: <u>bulik/ldsc: LD Score Regression (LDSC)</u> (github.com)
5. Two sample MR 0.4.26: <u>Two Sample MR functions and interface to MR Base database • TwoSampleMR</u> (mrcieu.github.io)
6. MR PResso: <u>GitHub - rondolab/MR-PRESSO: Performs the Mendelian Randomization Pleiotropy RESidual Sum and Outlier (MR-PRESSO) method</u>.
7. CAUSE: <u>Introduction to CAUSE</u> (jean997.github.io)
8. MR power calculator: <u>Online sample size and power calculator for Mendelian randomization with a binary outcome</u> (shinyapps.io)
9. MAGMA: https://ctg.cncr.nl/software/magma
10. HMAGMA: <u>GitHub - thewonlab/H-MAGMA</u>
11. GCTB: <u>GCTB</u> (cnsgenomics.com)
12. Polyfun (installed November 2021): <u>GitHub - omerwe/polyfun: PolyFun (POLYgenic FUNctionally-informed fine-mapping)</u>
13. SMR: <u>SMR | Yang Lab</u> (westlake.edu.cn)
14. Hyprcoloc: <u>GitHub - jrs95/hyprcoloc: Hypothesis prioritisation in multi-trait colocalization</u>
15. PRScs: <u>GitHub - getian107/PRScs: Polygenic prediction via continuous shrinkage priors</u>
16. R statistics packages: nlme, stats, Psychmeta, lmer, effsize, lsa, ape, factoExtra, igraph,
17. R plotting packages: ggseg, ggplot2, tidyverse, ggpubr, ggstatsplot, ComplexHeatmap, scico, karyoploteR, tidySEM, circlize

## Footnotes

1 We use the term “predominantly European genetic ancestries” to reflect ongoing conversations about terms used to refer to genetically homogenous groups. We recognise that the genetically homogenous group that clusters with European samples need not be entirely European or that their genetic ancestries are fully captured.

