## Supplemental information for "The genetics of cortical organisation and development: a study of 2,347 neuroimaging phenotypes"


**Supplementary Figures** 2

Supplementary Figure 1: Consistency in genetic effects between ABCD and UKB 2

Supplementary Figure 2: Clustering of global phenotypes using phenotypic (A) and genetic (B) correlation matrices. 3

Supplementary Figure 3: Phenotypic structural equation model 4

Supplementary Figure 4: Mendelian randomisation analysis for causal relationships between genetic effects on global brain phenotypes. 5

Supplementary Figure 5: Forest plots and leave-one-out plots 6

Supplementary Figure 6: Co-localisation lots of all clusters with cluster size > 30 7

Supplementary Figure 7: Co-localisation lots of all clusters with cluster size > 30 8

Supplementary Figure 8: Co-localisation and regional association plot for FI and ICI-6:125424383-127540461 9

Supplementary Figure 9: Regional heritability 10

Supplementary Figure 10: Spin permutation based enrichment of mean SNP heritability across mesulam classes and Yeo & Krienen communities. 11

Supplementary Figure 11: Spin permutation based enrichment of mean genetic correlations within mesulam classes and Yeo & Krienen communities. 12

Supplementary Figure 12: Topography of the first phenotypic principal components. 13

**Supplementary notes and associated figures** 14

Supplementary Note 1: Phenotypic factor analysis and structural equation modelling 14

Supplementary Note 2: Identifying causal factors for folding using Mendelian Randomisation 15

Supplementary Note 3: Adjusting for global phenotypes can bias regional GWAS 16

Supplementary Figure 13: Acyclic graphs for the association between genetic variant and regional phenotypes 17

Supplementary Figure 14: Correlation between regional phenotypes and global phenotypes 18

**References** 19

### Supplementary Figures

#### Supplementary Figure 1: Consistency in genetic effects between ABCD and UKB

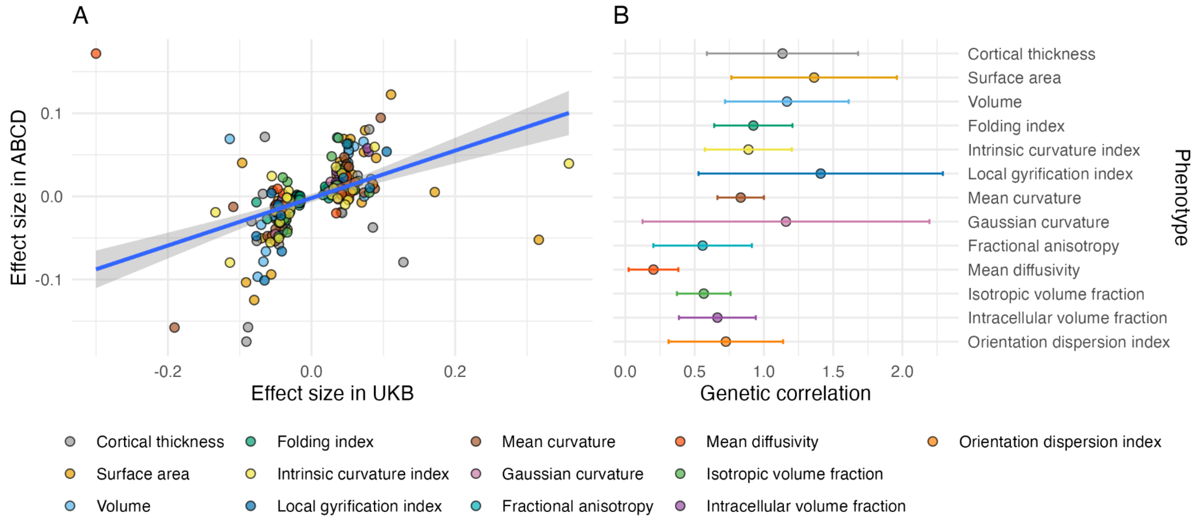


*(A) Correlation in effect size (regression beta from GWAS) between ABCD and UKB cohorts for all 237 genome-wide significant SNPs in the UKB: Pearson's correlation coefficient, r =0.54 with 95% confidence interval 0.45-0.63. (B) Genetic correlation (points) and 95% confidence intervals (lines) for 12 global phenotypes in the UKB and ABCD cohorts. Given the relatively small size of ABCD, the intercept has been constrained as there is no participant overlap between the UKB and ABCD and there is no inflation in test statistics due to uncontrolled population stratification. Consequently, estimates of genetic correlation can be above 1.*

Supplementary Figure 2: Clustering of global phenotypes using phenotypic (A) and genetic (B) correlation matrices.

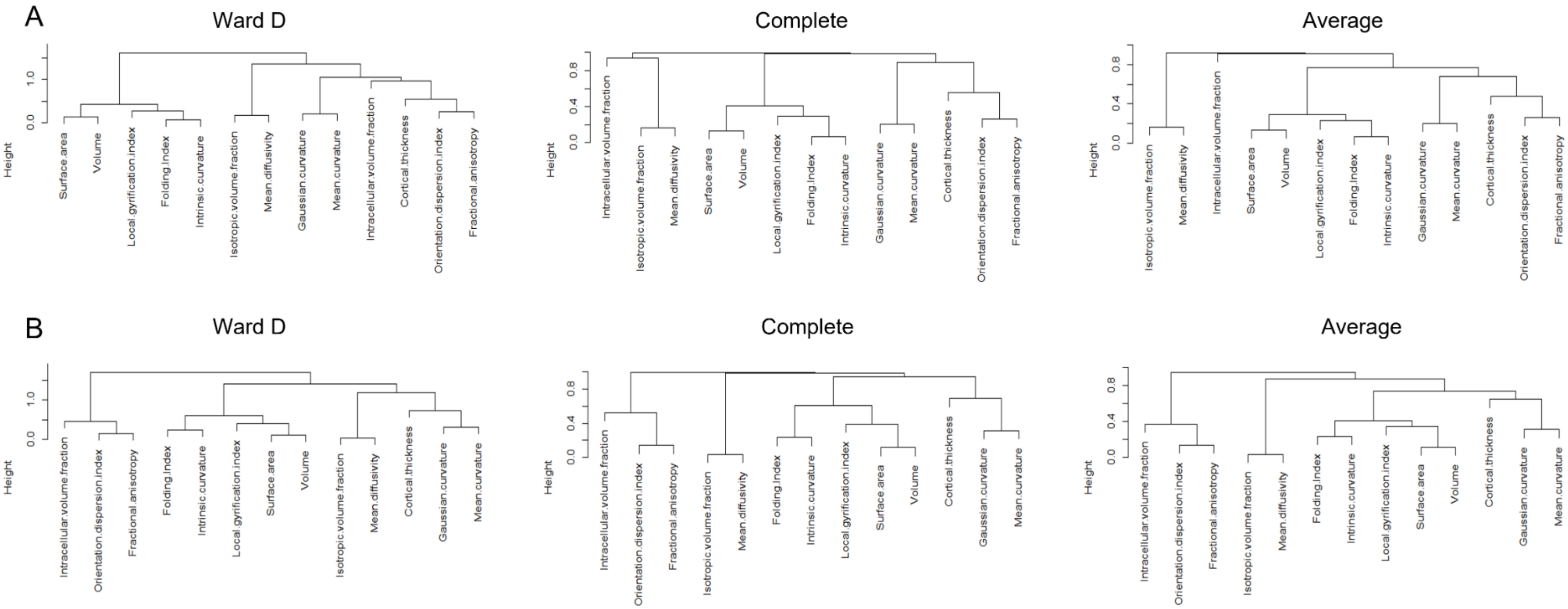


*Consistency of clustering approaches was assessed using 3 methods for hierarchical clustering: clustering using Ward's criterion, complete linkage, and average linkage.*

Supplementary Figure 3: Phenotypic structural equation model

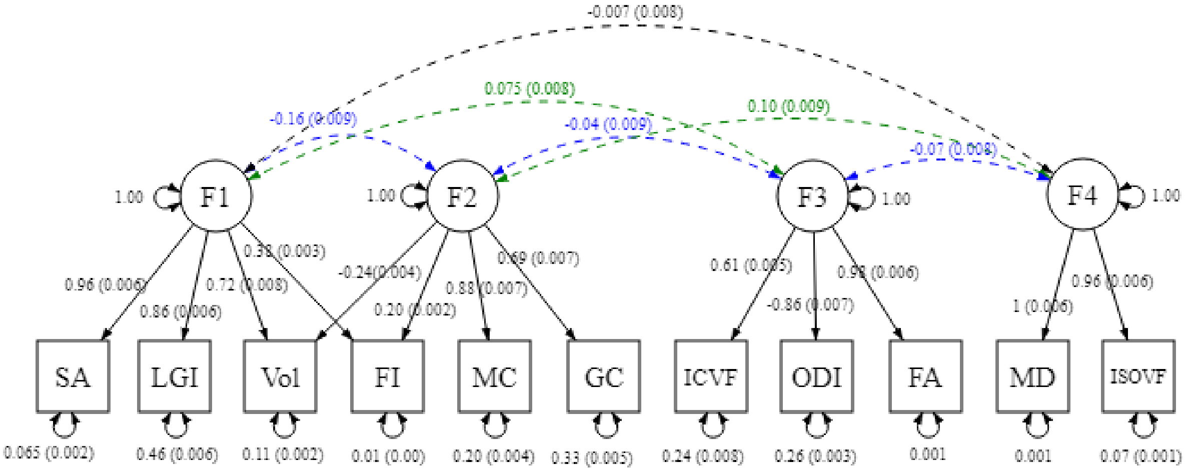


*Phenotypic SEM path diagram demonstrating the underlying latent structure of 12 of the 13 global phenotypes and the interfactor genetic correlations. Covariance relationships = double-headed arrows connecting two variables, variance estimates = double-headed arrows connecting variable to itself, regression relationships = one-headed arrows pointing from independent variable to dependent variable. Circles indicate latent variables, squares indicate measured phenotypes. Abbreviations: cortical surface area (SA), grey matter volume (Vol), folding index (FI), local gyrification index (LGI), mean curvature (MC), gaussian curvature (GC), fractional anisotropy (FA), mean diffusivity (MD), intracellular volume fraction (ICVF), isotropic volume fraction (ISOVF), and orientation dispersion index (ODI).*

Supplementary Figure 4: Mendelian randomisation analysis for causal relationships between genetic effects on global brain phenotypes.

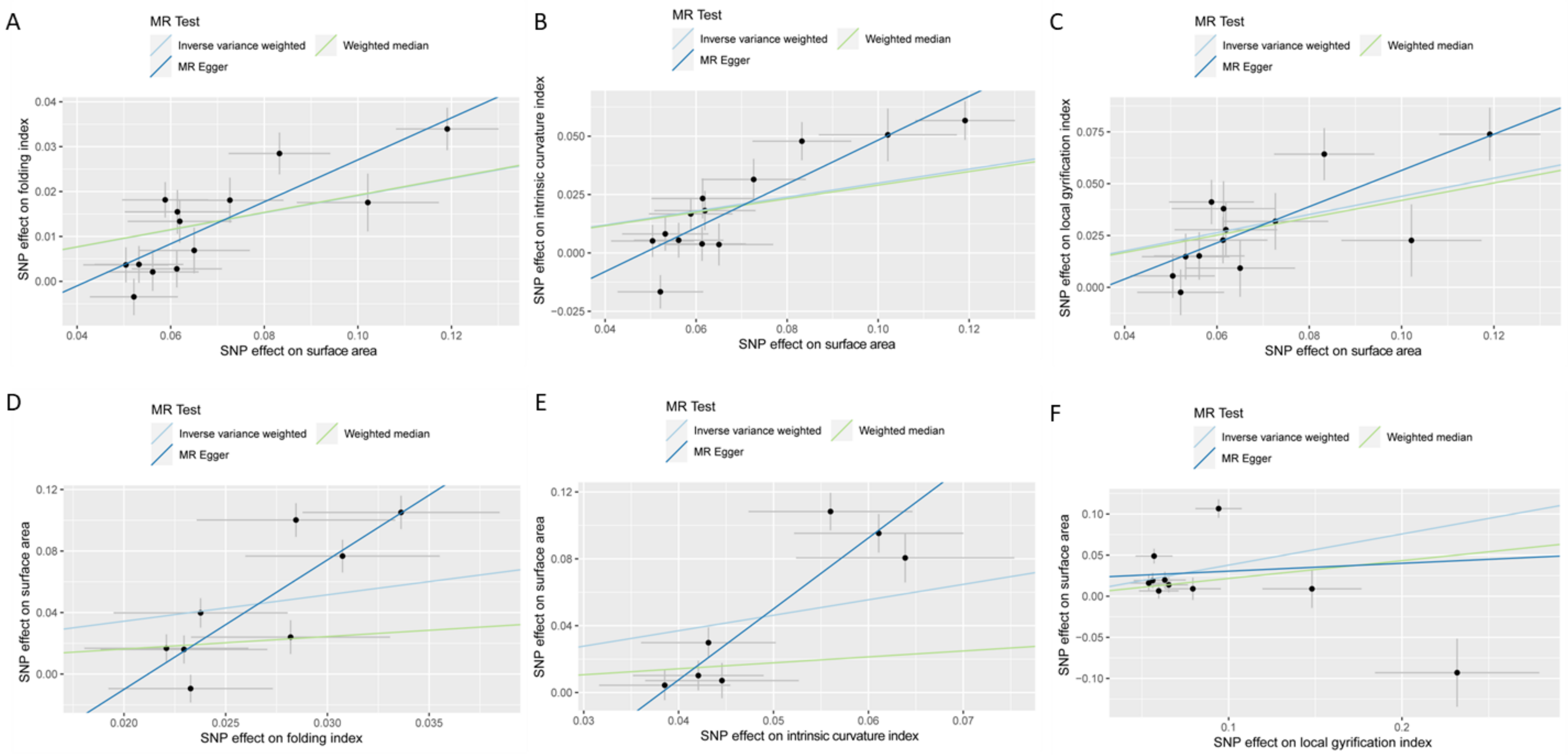


*Scatter plots for the bidirectional MR effects between surface area and folding index, intrinsic curvature index, and local gyrification index. A, B, and C are scatterplots where surface area is the exposure, and D, E, and F are scatterplots where surface area is the outcome. All scatter plots are for MR analyses conducted by splitting the UKB into two samples of similar sample sizes. All estimates were statistically significant in scatterplots A,B, and C. Inverse-variance weighted MR failed to reach statistical significance in scatterplots D,E, and F.*

Supplementary Figure 5: Forest plots and leave-one-out plots

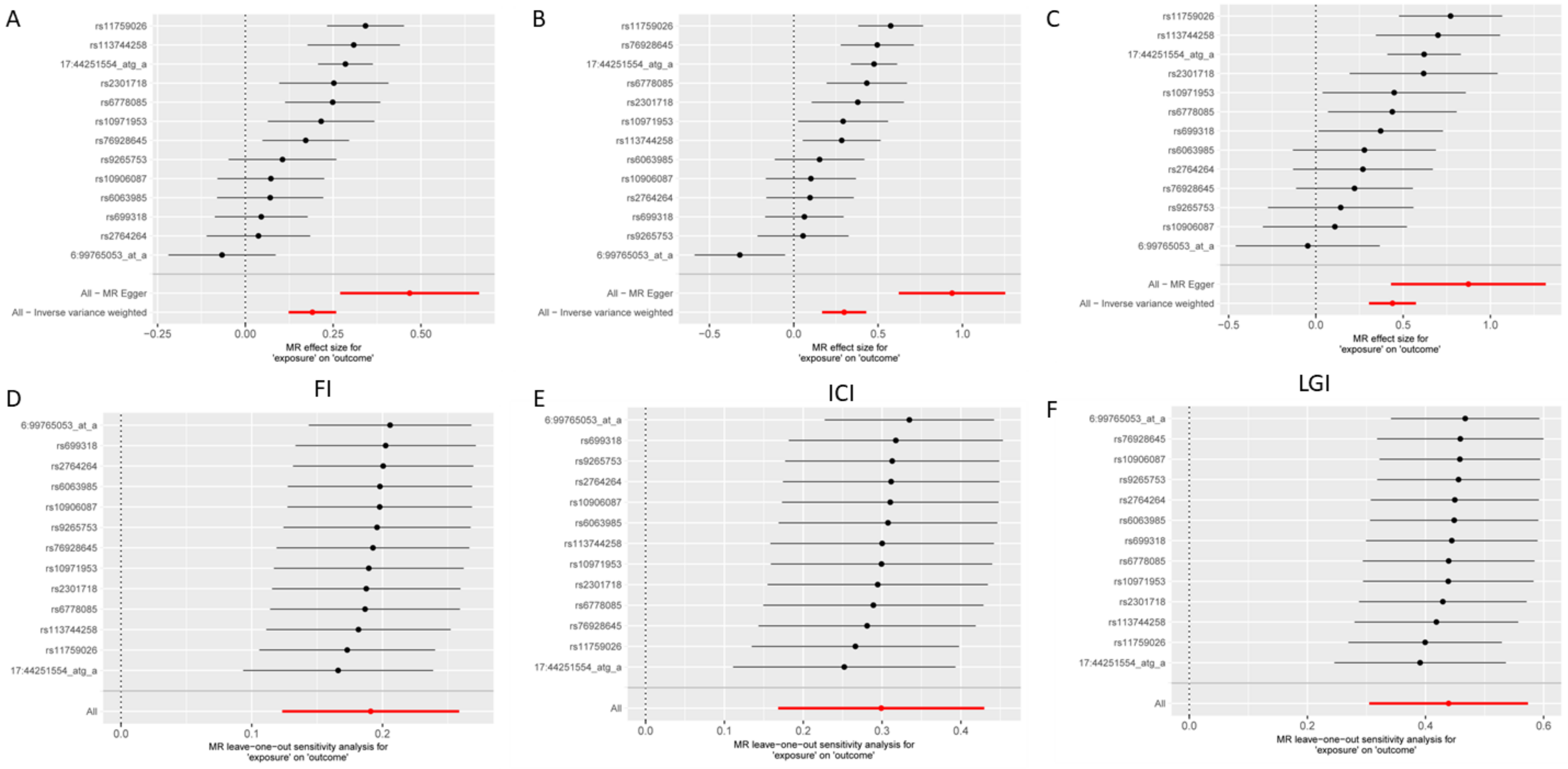


Forest plots (A, B, and C) and leave-one-out (D, E, and F) between surface area and folding index (FI, A and D), Intrinsic curvature index (ICI, B and E), and local gyrification index (LGI, C and F).

Supplementary Figure 6: Co-localisation lots of all clusters with cluster size > 30

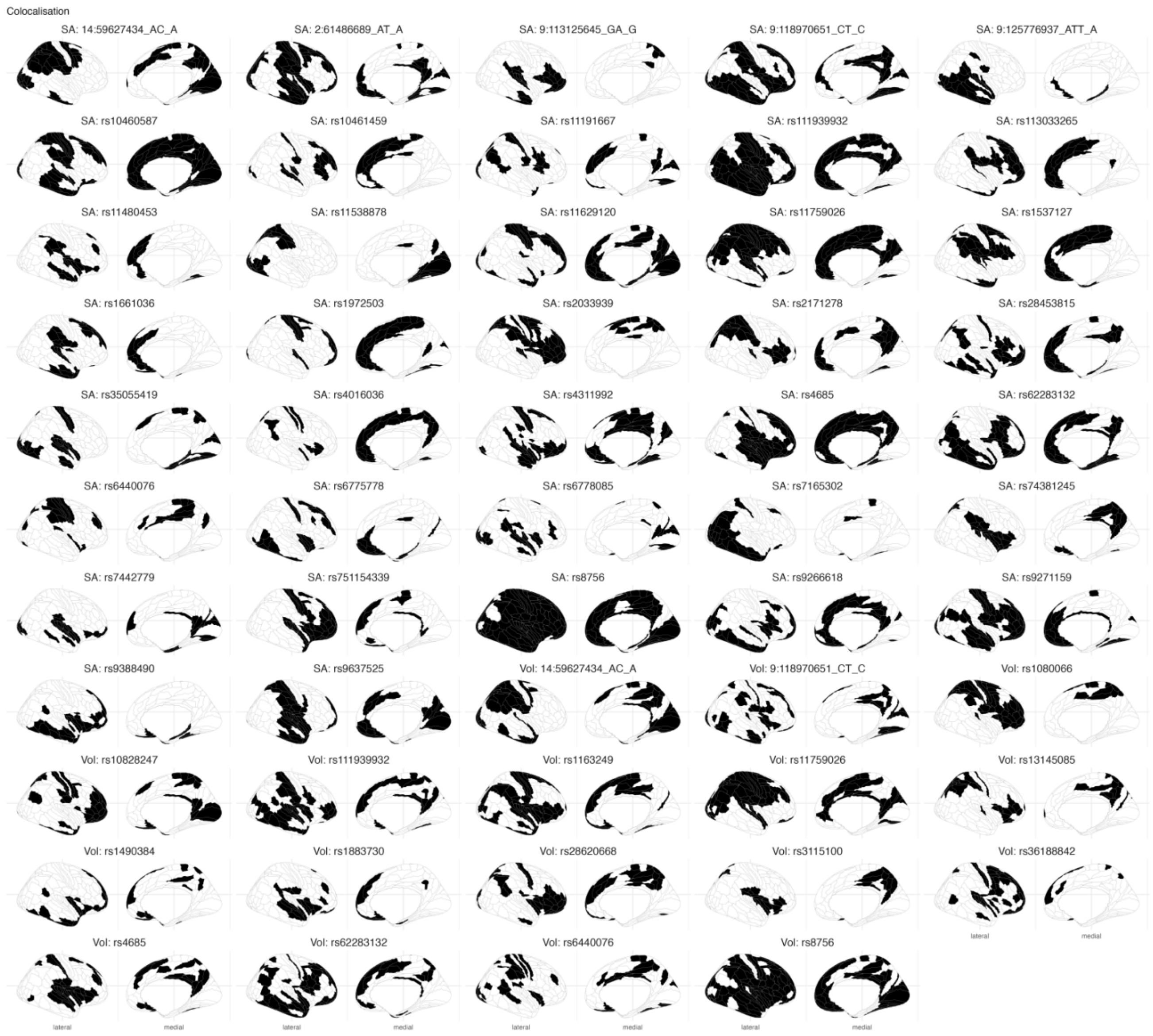


Cortical topographical plots demonstrating topographical distribution of co-localisation clusters with cluster size 30 for volume and surface area. Clusters are coloured in black. For each plot, the relevant phenotype and the SNP identified as the candidate causal variant for the cluster is provided.

Supplementary Figure 7: Co-localisation lots of all clusters with cluster size > 30

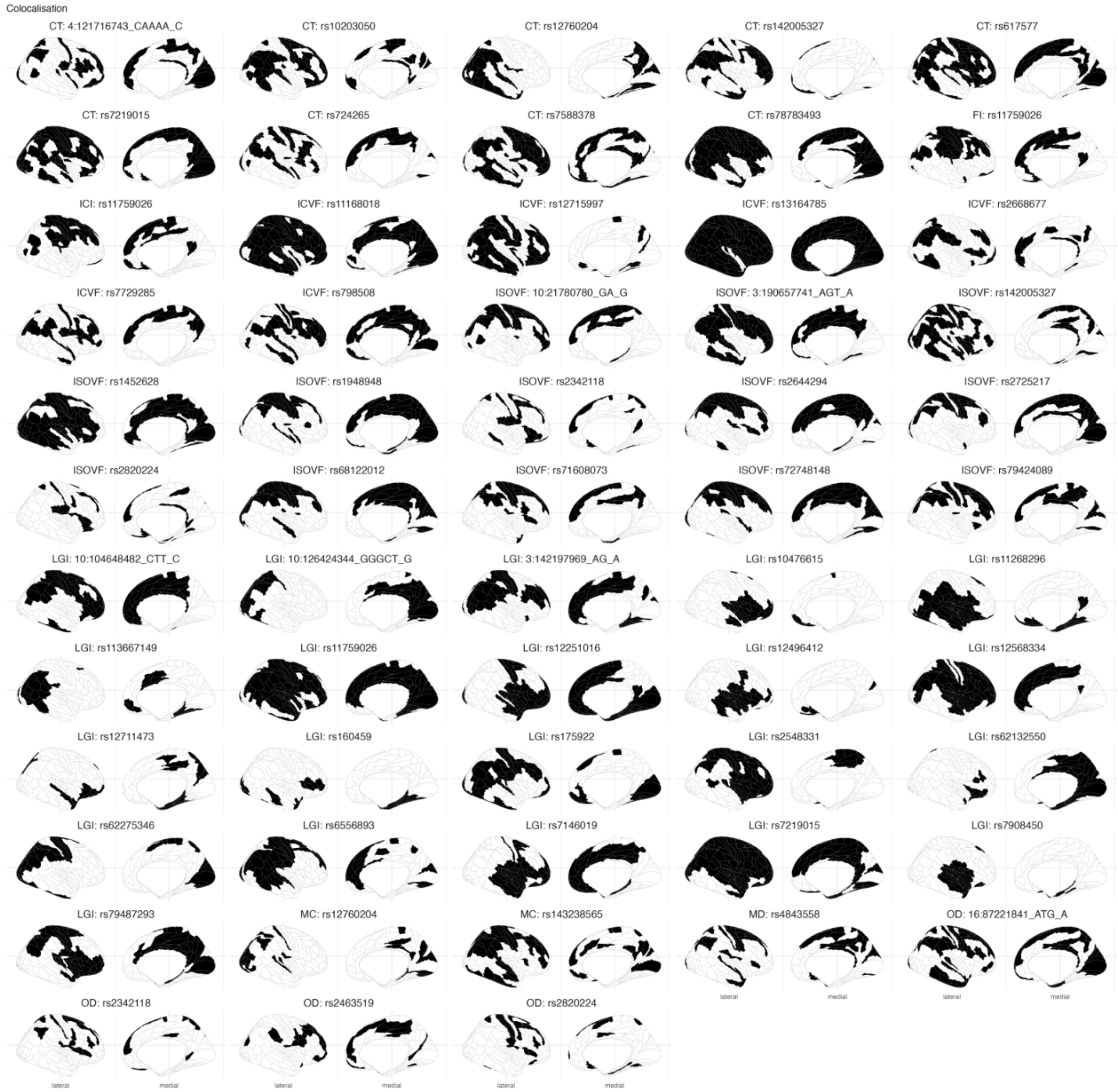


Cortical topographical plots demonstrating topographical distribution of co-localisation clusters with cluster size 30 for all phenotypes except volume and surface area. Clusters are coloured in black. For each plot, the relevant phenotype and the SNP identified as the candidate causal variant for the cluster is provided.

Supplementary Figure 8: Co-localisation and regional association plot for FI and ICI-6:125424383-127540461

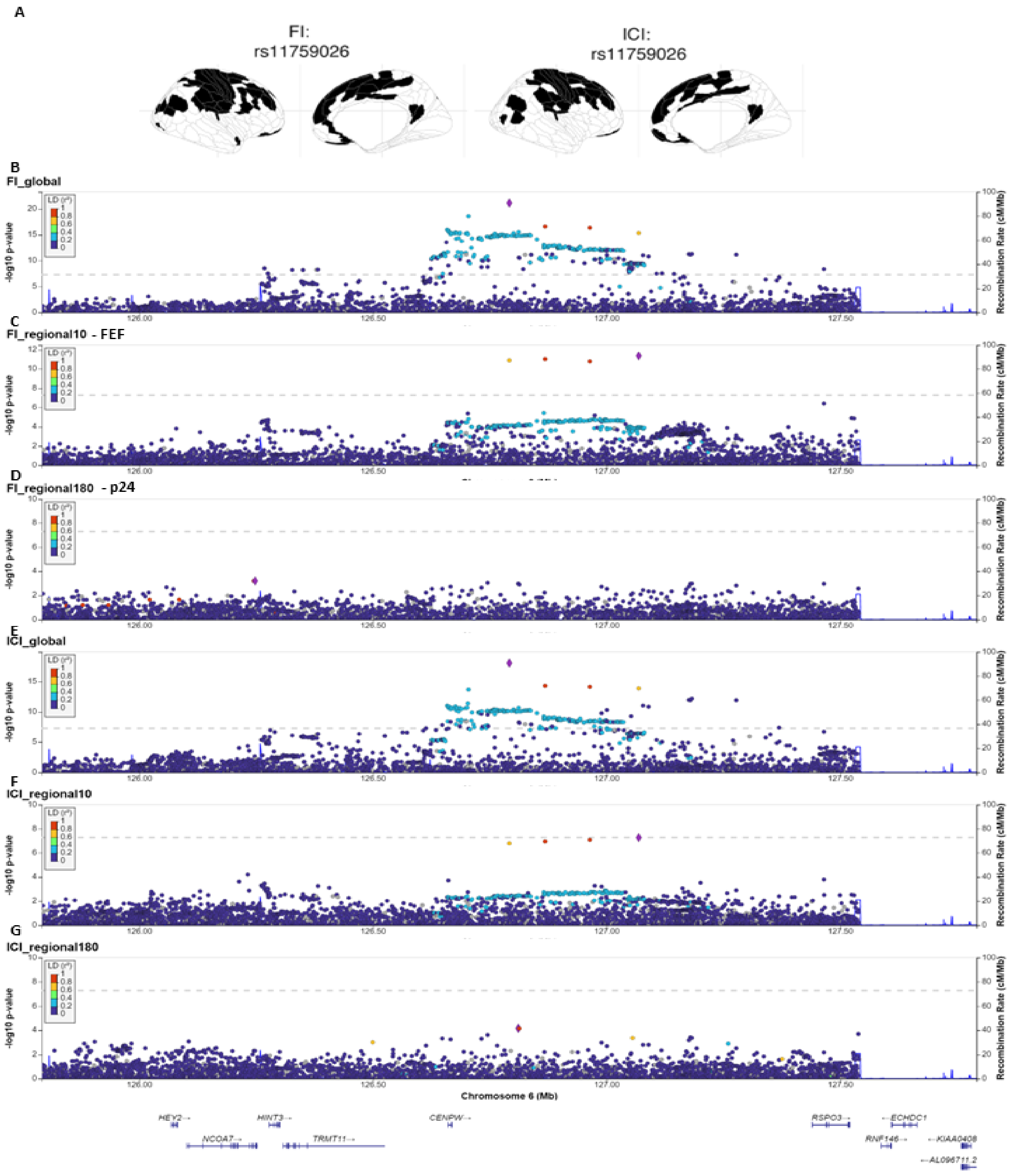


A: Cortical topographic plots of two co-localised loci with ICI and FI. B and E: Global GWAS for FI and ICI respectively. C and F regional GWAS (FEF region) that was part of the co-localised cluster in FI and ICI respectively. D and G: regional GWAS (p24 region) that was outside the co-localised cluster in FI and ICI.

Supplementary Figure 9: Regional heritability

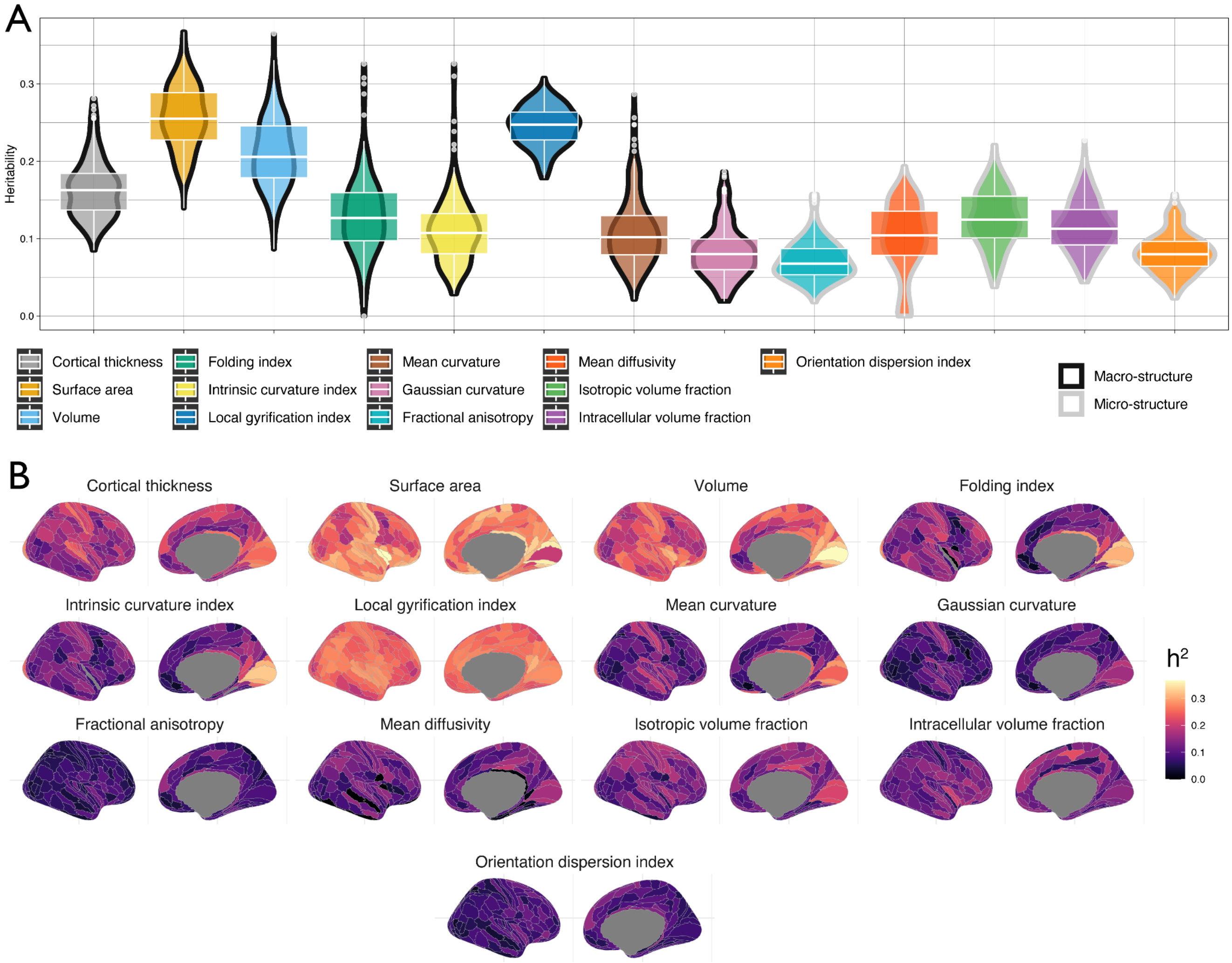


A. The distribution of the SNP heritability for the regional phenotypes of the 13 neuroimaging modalities. B. The cortical spatial topology of SNP heritability for the 13 neuroimaging modalities.

Supplementary Figure 10: Spin permutation based enrichment of mean SNP heritability across mesulam classes and Yeo & Krienen communities.

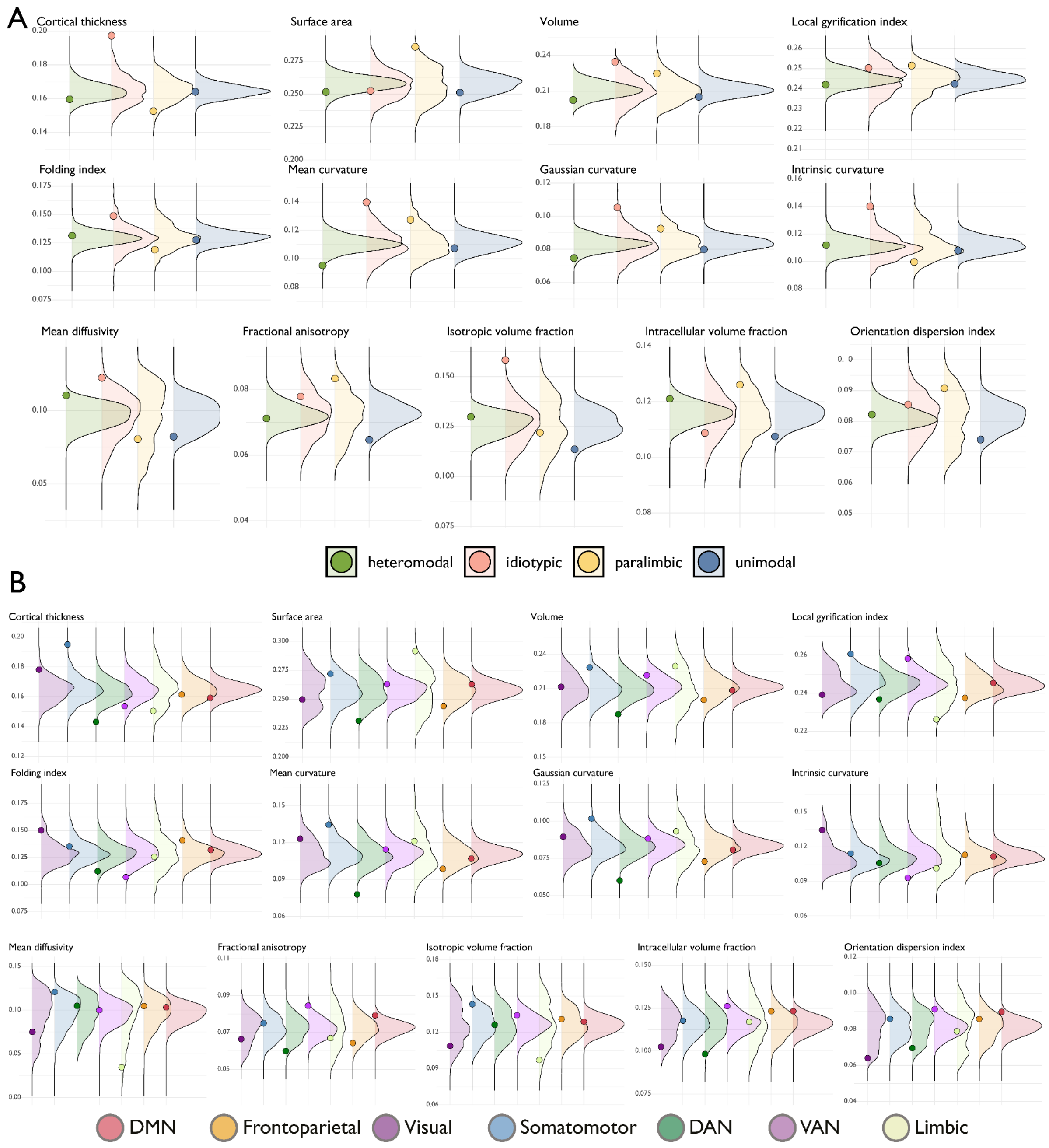


A. The average SNP heritability for each Mesulam class (dots) overlaid onto a null distribution of the heritability in that class obtained by spinning the parcellation 1000 times (and computing the average heritability within each permutation). Thus, if the real average heritability (dots) are within the tail ends (5%) of the null distribution it could be concluded that this heritability is higher in that class and for that feature than would be expected from a spatially random distribution of the heritability across the cortex. Only idiotypic regions for some phenotypes (cortical thickness, volume, mean, gaussian and intrinsic curvature, mean diffusivity and isotropic volume fraction) show this relative spatial enrichment. B. The same type of spatial enrichment analysis for the Yeo and Krienen communities.

Supplementary Figure 11: Spin permutation based enrichment of mean genetic correlations within mesulam classes and Yeo & Krienen communities.

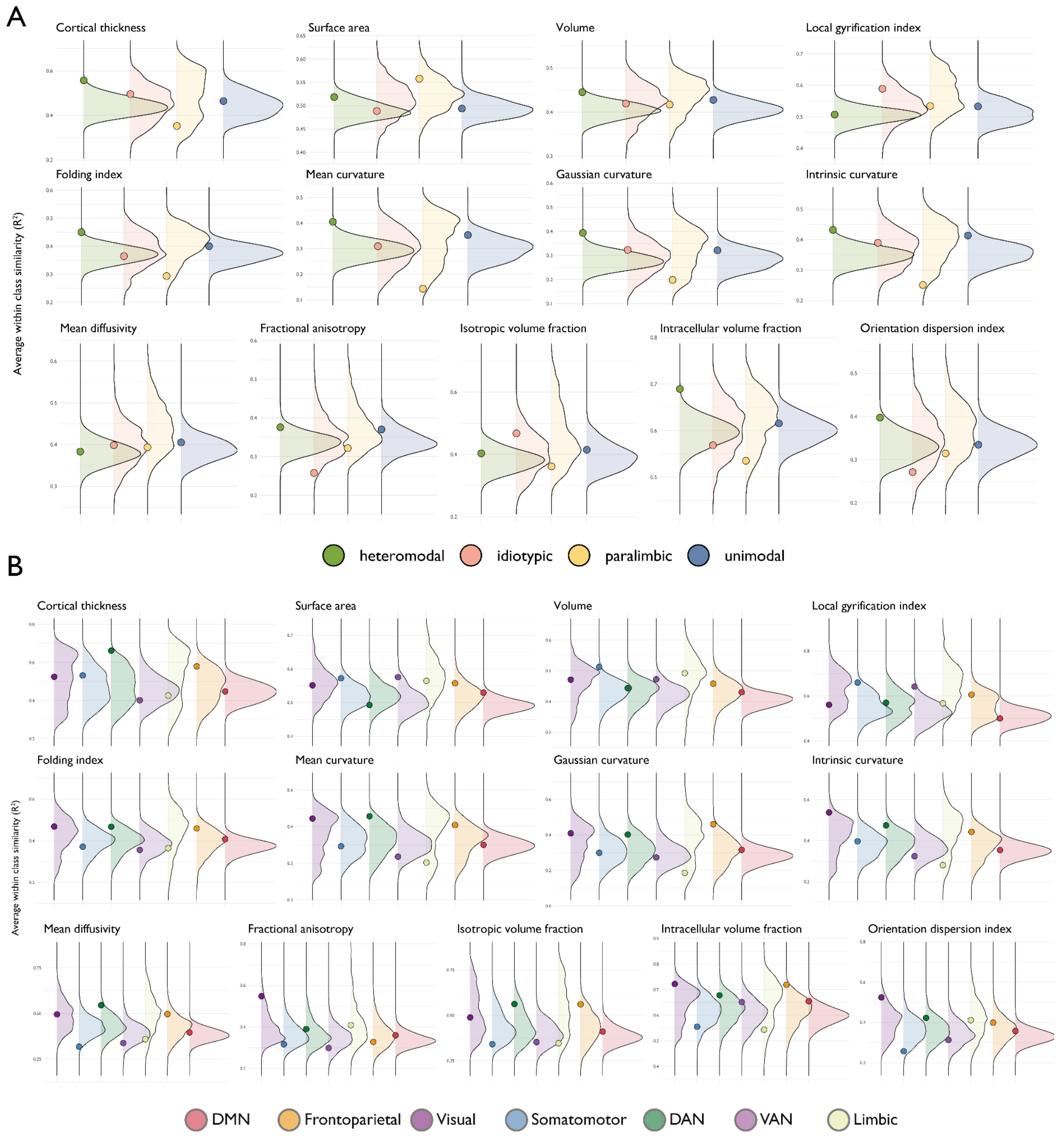


A. The average within class genetic similarity (i.e., the average of all edges in the genetic correlation matrix for regions belonging to the same class) for each Mesulam class (dots) overlaid onto a null distribution obtained by spinning the parcellation 1000 times (and computing the average similarity within each permutation). Thus, if the real genetic correlations for a given class (dots) are within the tail ends (5%) of the null distribution it could be concluded that this genetic correlation within regions belonging to the same class is higher in that class and for that feature than would be expected from a spatially random distribution across the cortex. B. The same type of spatial enrichment analysis for the Yeo and Krienen communities.

Supplementary Figure 12: Topography of the first phenotypic principal components.

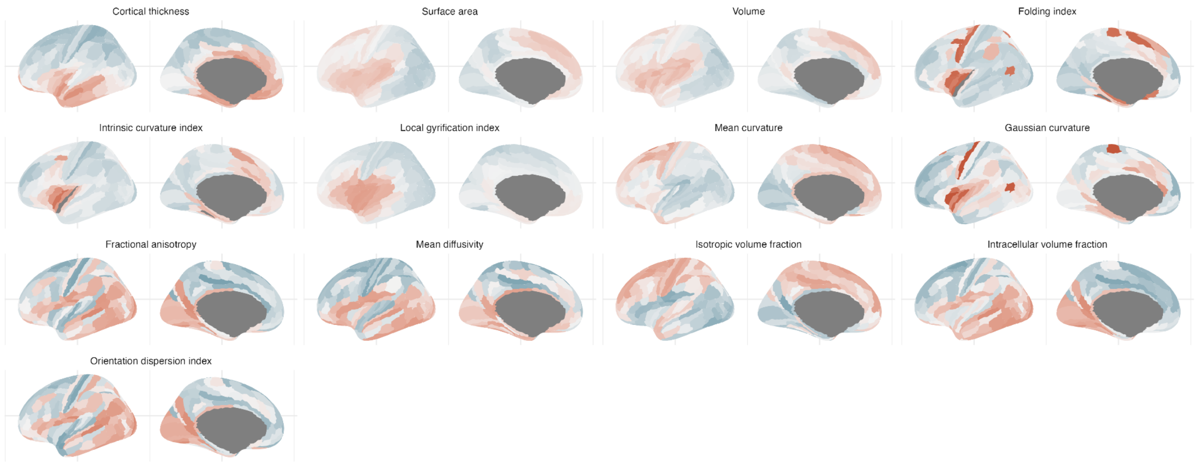


Colorscales depict the relative eigenvector ranging from -20 to +29, in all plots the midpoint is defined as 0. It should be notes that the definition of $Ax=\lambda x$ as the PCA eigenvector means that the sign is somewhat ambiguous and that the magnitude is relative to its own scaling (in this case within each phenotype for which the PCA is performed). Thus, in this context, the colour scale indicates to what extent regions show more homogenous similarity (i.e., regions with more similar colour have more similar covariance).

#### Supplementary notes and associated figures

##### Supplementary Note 1: Phenotypic factor analysis and structural equation modelling

Using the quality controlled phenotypic dataset which was included in the GWAS, we conducted exploratory factor analysis on half of the sample ($N_{total}$ = 28,794, $N_{half}$ = 14,397 ), randomly selected. We conducted confirmatory factor analysis in the other half of the sample. All phenotypes were scaled. Examination of the phenotypic correlation matrix indicated that, with the exception of CT, most phenotypes exhibited a pattern of high absolute correlation with at least one other phenotype and low correlations with others. In contrast, CT exhibited moderate correlation with almost all phenotypes, in line with the idea that multiple cell types underlie CT.

We next investigated if our data is amenable to factor analysis using Bartlett's test of sphericity Kaiser-Meyer-Olkin measure of sampling adequacy. The overall measure of sampling adequacy was 0.5, which is the minimum acceptable for factor analysis. Inspecting individual measures of sampling adequacy identified very low sampling adequacy for CT (0.29), consistent with earlier findings that CT does not cluster very well with other phenotypes. Excluding CT produced a better overall measure of sampling adequacy (0.57). Bartlett's test confidence was significant both before and after excluding CT.

Given the above points, we excluded CT from factor analysis and conducted exploratory factor analysis on 12 of the 13 global phenotypes. Scree plot and parallel analysis indicated four factors, in line with the number of clusters identified from hierarchical clustering. Exploratory factor analysis identified four factors: 1: Cortical expansion; 2: Cortical curvature; 3: Neurite density; and 4: Water diffusion (**SF 3**). There was significant and high (> 0.5) cross-loading of ICI onto two factors (cortical expansion and cortical curvature). Multiple iterations of the confirmatory factor analyses when including ICI failed to produce satisfactory models. After removing ICI, we identified a similar four factor model relating to cortical expansion, cortical curvature, neurite density and water diffusion with acceptable fit indices (CFI: 0.85, SRMR: 0.79).

#### Supplementary Note 2: Identifying causal factors for folding using Mendelian Randomisation

Understanding the processes that underlie cortical folding has proven elusive. Several theories have tried to explain folding including skull constraint, axonal tension, and differential tangential growth[1]. Skull constraint and axonal tension do not account for the uniform nature of folding across among humans. The cellular model for differential tangential expansion posits that folding occurs through two mechanisms: first, the outer layers expand more than the inner layers, causing the cortex to fold and second, there is heterogeneous distribution of progenitors across the brains, leading to differences in neurogenesis[1–3]. In humans, one way to test this is to investigate if genetic variants underlying surface area increase measures of curvature. We used multiple Mendelian Randomisation methods to investigate this.

First, we split the UK Biobank into two datasets of roughly equal sample size to ensure that there is limited participant overlap and conducted MR. Inverse-variance weighted MR demonstrated that genetically predicted surface area increased genetically predicted local gyrification index (LGI), intrinsic curvature index (ICI), and folding index (FI) after Bonferroni correction. These results were statistically significant after applying methods that are robust to various assumptions: median-weighted[4] MR and MR-Presso[5], and after removing outliers by Steiger filtering[6]. These results also had consistent effect direction when using MR-Egger[7], which is statistically underpowered compared to other methods. However, in the reverse direction, we did not obtain consistent results to indicate that genetic variants associated with these measures increased genetically predicted surface area (**SF 4**). Visual inspection of the forest plots and leave-one-out plots did not indicate that the results were driven by one or two genetic variants (**SF 5**).

We confirmed our findings first by using a different method that can model both correlated and uncorrelated pleiotropy (CAUSE)[8] using two sets of instruments - one created from SNPs with $p < 5x10^{-8}$, and another with SNPs with $p < 1x10^{-3}$, as previously demonstrated by the developers of the method. CAUSE suggested that SA causally increased LGI, FI and ICI. In the reverse direction, CAUSE identified a shared model between LGI, FI, and SA, and a causal model between SA and ICI. Consistent results suggesting that genetically predicted surface area increased genetically predicted measures of folding were obtained when MR was run using ABCD and UKB. Altogether, MR provides support for the differential tangential growth theory for some measures of folding, but suggests that other mechanisms may contribute to other measures of folding.

#### Supplementary Note 3: Adjusting for global phenotypes can bias regional GWAS

Following Aschard and colleagues[9], we consider four scenarios in which a genetic variant may be associated with regional phenotypes. Acyclic graphs are provided below (**SF 13**).

Scenario 1: A genetic variant is associated with a global phenotype but not associated with a regional phenotype.

Scenario 2: A genetic variant is associated with a regional phenotype but not associated with global phenotypes.

Scenario 3: A genetic variant is independently associated with both regional and global phenotypes.

Scenario 4: The genetic effect of a variant on a regional phenotype is mediated partly or completely by global phenotypes.

For Scenarios 2 and 4, controlling for the global phenotype will not affect the association between the genetic variant and the regional phenotype. However, for scenarios 1 and 3, correcting for the global phenotype will induce a correlation between the genetic variant and the regional phenotype. The estimate will be biased by $\beta_C\rho_{CY}$ where $\beta_C$ is the effect of the genetic variant on the covariate (in this case the global phenotype), and $\rho_{CY}$ is the correlation between the covariate (the global phenotype) and outcome. In our study, the $\beta_C$ for the standardised global phenotypes range from (-1.47) to (1.16) for the genome-wide significant loci, and the correlations between the global phenotypes and regional phenotypes are high and displayed in (**SF 14**). Therefore, correcting for the global phenotypes will bias the regional estimates.

Supplementary Figure 13: Acyclic graphs for the association between genetic variant and regional phenotypes

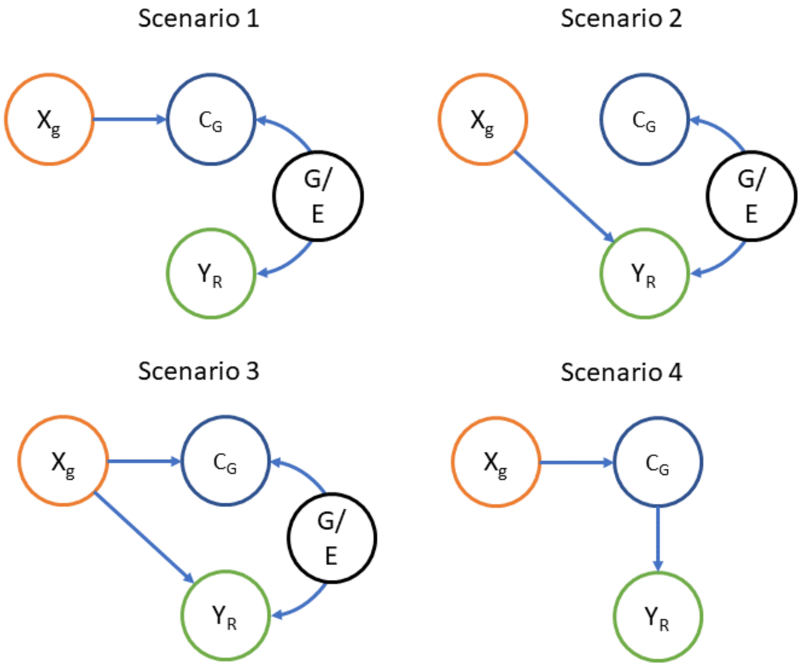


Acyclic graphs demonstrating causal relationship between genetic variant ($X_g$) and regional phenotype ($Y_R$), in the presence of the global phenotype which is the covariate ($C_G$) and other genetic and environmental factors (G/E) which contribute to the phenotypic correlation between the regional phenotype and the global phenotype.

Supplementary Figure 14: Correlation between regional phenotypes and global phenotypes

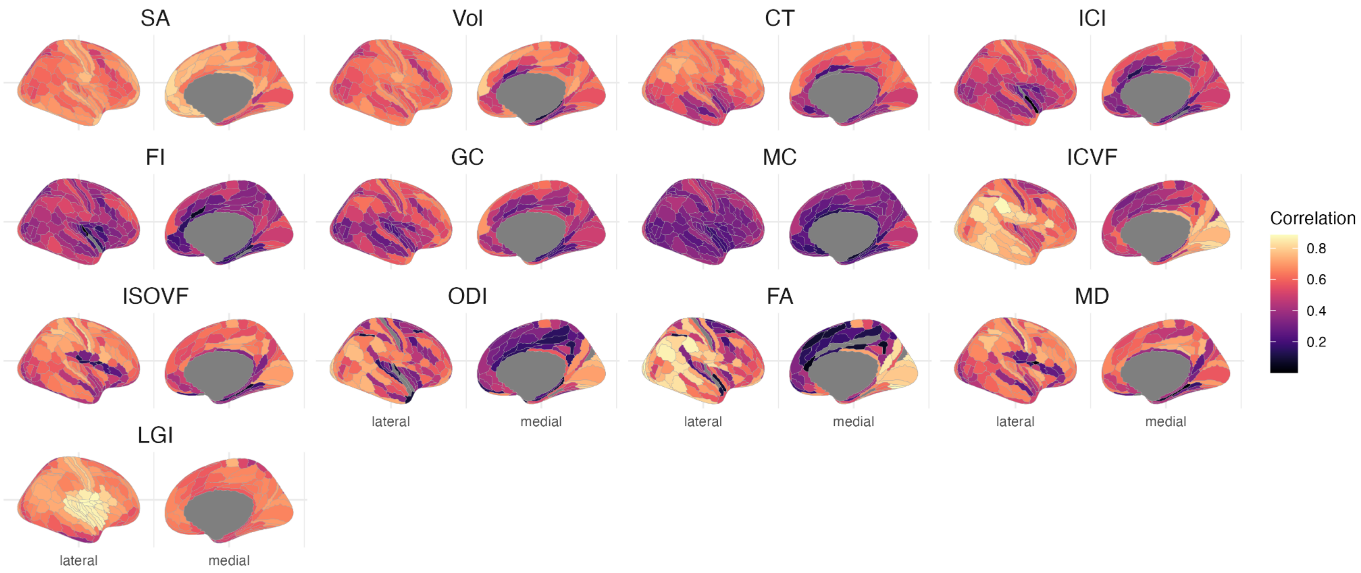


Pearson’s correlation coefficient between regional phenotypes and global phenotypes.
